# Mean-field approximations with adaptive coupling for networks with spike-timing-dependent plasticity

**DOI:** 10.1101/2022.07.02.498537

**Authors:** Benoit Duchet, Christian Bick, Áine Byrne

## Abstract

Understanding the effect of spike-timing-dependent plasticity (STDP) is key to elucidate how neural networks change over long timescales and to design interventions aimed at modulating such networks in neurological disorders. However, progress is restricted by the significant computational cost associated with simulating neural network models with STDP, and by the lack of low-dimensional description that could provide analytical insights. Phase-difference-dependent plasticity (PDDP) rules approximate STDP in phase oscillator networks, which prescribe synaptic changes based on phase differences of neuron pairs rather than differences in spike timing. Here we construct mean-field approximations for phase oscillator networks with STDP to describe part of the phase space for this very high dimensional system. We first show that single-harmonic PDDP rules can approximate a simple form of symmetric STDP, while multi-harmonic rules are required to accurately approximate causal STDP. We then derive exact expressions for the evolution of the average PDDP coupling weight in terms of network synchrony. For adaptive networks of Kuramoto oscillators that form clusters, we formulate a family of low-dimensional descriptions based on the mean field dynamics of each cluster and average coupling weights between and within clusters. Finally, we show that such a two-cluster mean-field model can be fitted to synthetic data to provide a low-dimensional approximation of a full adaptive network with symmetric STDP. Our framework represents a step towards a low-dimensional description of adaptive networks with STDP, and could for example inform the development of new therapies aimed at maximizing the long-lasting effects of brain stimulation.

## 1 Introduction

Synaptic plasticity is considered the primary mechanism for learning and memory consolidation. Neurons with similar activity patterns strengthen their synaptic connections, while others connections may weaken. Spike-timing-dependent plasticity (STDP) has been suggested as an unsupervised, local learning rule in neural networks (Gerstner, Kempter, Van Hemmen, & Wagner, 1996; Song, Miller, & Abbott, 2000) motivated by experimental findings (Markram, Lübke, Frotscher, & Sakmann, 1997; Bi & Poo, 1998; Feldman, 2000; Froemke & Dan, 2002; Cassenaer & Laurent, 2007; Sgritta, Locatelli, Soda, Prestori, & D’ Angelo, 2017). These experimental studies reported synaptic strengthening or weakening depending on the order and timing of pre- and postsynaptic spikes. In causal STDP (Fig. 1A), long-term potentiation (LTP) occurs when the postsynaptic neuron fires shortly after the presynaptic neuron. Conversely, long-term depression (LTD) occurs when the postsynaptic neuron fires shortly before the presynaptic neuron. The closer the spike times are, the larger the effect. Non-causal, symmetric STDP has also been reported in the hippocampus (Abbott & Nelson, 2000; Mishra, Kim, Guzman, & Jonas, 2016), see Fig. 1B. Although Donald Hebb emphasized the importance of causality in his theory of adaptive synaptic connections in 1949 (Hebb, 2005), such non-causal rules, that neglect temporal precedence, are sometimes referred to as Hebbian plasticity rules.^1^

**Figure 1:**
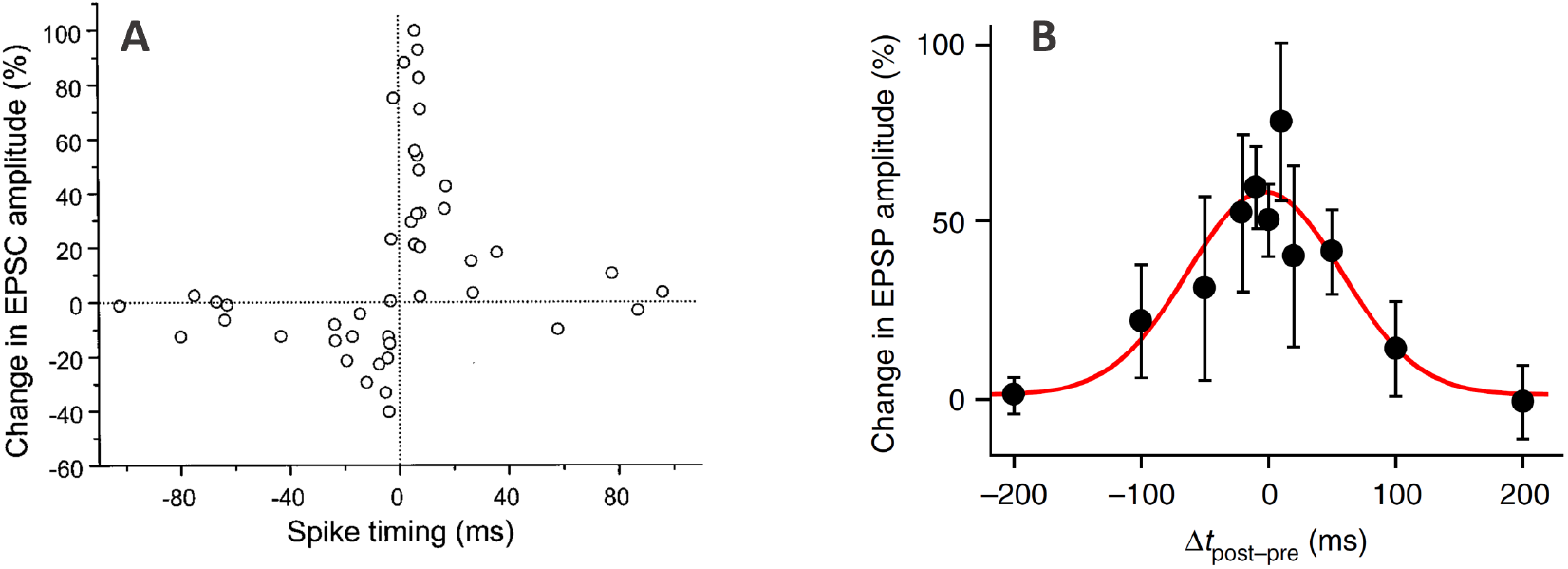
Examples of STDP observed in pairing experiments. In typical STDP pairing experiments, the presynaptic neuron is stimulated shortly before or after forcing the postsynaptic neuron to fire by injecting a brief current pulse (with a controlled delay). The pairing is repeated many times for each value of the delay. **A:** causal STDP in synapses on glutamatergic neurons in rat hippocampal culture. Adapted from (Bi & Poo, 1998) with no permission required (Copyright 1998 Society for Neuroscience). **B:** symmetric STDP in CA3-CA3 synapses in the hippocampus (slices in the rat). Adapted from (Mishra et al., 2016) with no permission required. In both panels, the horizontal axes represent the difference in spike timing (postsynaptic minus presynaptic), and the vertical axes represent measures of the change in synaptic strength, either involving excitatory postsynaptic currents (EPSC, panel A), or excitatory postsynaptic potentials (EPSP, panel B).

As a major contributor to long-term plasticity (time scale of 1s or longer), STDP is key to long-term neural processes in the healthy brain, such as memory (Litwin-Kumar & Doiron, 2014) and sensory encoding (Coulon, Beslon, & Soula, 2011), but also to modulate networks affected by neurological disorders (Madadi Asl, Vahabie, Valizadeh, & Tass, 2022). In particular, long-term plasticity is critical to the design of effective therapies for neurological disorders based on invasive and non-invasive brain stimulation. Paired associative stimulation using transcranial magnetic stimulation (TMS) has been shown to trigger STDP-like changes (Müller-Dahlhaus, Ziemann, & Classen, 2010; Johnen et al., 2015; Wiratman et al., 2022), and STDP models have been used to design electrical stimulation for stroke rehabilitation (Kim et al., 2021). Coordinated reset deep brain stimulation (DBS) for Parkinson’s disease was designed to induce long-term plastic changes outlasting stimulation using STDP models (Tass & Majtanik, 2006), and was later validated in non-human primates (Tass et al., 2012; Wang et al., 2016) and patients (Adamchic et al., 2014).

However, progress in these areas is restricted by the significant computational cost associated with simulating neural network models with STDP and by the lack of lowdimensional description of such networks. Indeed, simulating networks with STDP of even moderate size for more than a couple of minutes of biological time can be impractical because the number of weight updates scales with the square of the network size. Low-dimensional mean-field approximations have been developed for networks with short-term plasticity (Tsodyks, Pawelzik, & Markram, 1998; Taher, Torcini, & Olmi, 2020; Gast, Schmidt, & Knösche, 2020; Schmutz, Gerstner, & Schwalger, 2020; Gast, Knösche, & Schmidt, 2021), but reductions for STDP have assumed that plasticity does not change firing rates and spike covariances (Ocker, Litwin-Kumar, & Doiron, 2015) or that the network is in a balanced state at every point in time (Akil, Rosenbaum, & Josić, 2021). Even if node dynamics are given by a simple Kuramoto-type model, it is a challenge to understand the dynamics of large networks with adaptivity. While the continuum limit of Kuramoto-type networks with STDP-like adaptivity can be described by integro-differential equations from a theoretical perspective (Gkogkas, Kuehn, & Xu, 2021), these do not necessarily elucidate the resulting network dynamics or yield a computational advantage. Approaches like the Ott–Antonsen reduction (Ott & Antonsen, 2008), which have been instrumental to derive low-dimensional descriptions of phase oscillators (see (Bick, Goodfellow, Laing, & Martens, 2020) and references therein) are not directly applicable to adaptive networks where all connection weights evolve independently of one another. Indeed, one would not expect an exact mean-field description with only a few degrees of freedom to be possible (as for the Kuramoto model with static connectivity) without making further assumptions, as any exact low-dimensional description would have to reflect the high, adaptivity-induced multistability (Berner, Schöll, & Yanchuk, 2019).

Models of causal, additive STDP based on differences in spike timing are exemplified by the work of Song, Miller and Abbott (Song et al., 2000). They assumed that the synaptic strength *κ_kl_* from presynaptic neuron *l* to postsynaptic neuron *k* is updated only when either neurons spike, and the corresponding change in synaptic strength from *l* to *k* is given by

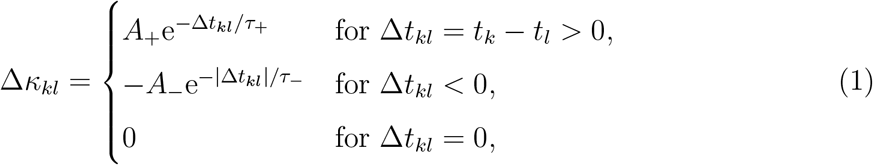

where the most recent spike times of neurons *k* and *l* are *t_k_* and *t_l_*, the parameters *A*_+_ and *A*_ determine the magnitude of LTP and LTD, and *τ*_+_ and *τ*_ determine the timescale of LTP and LTD, respectively (see Fig. 4A). Conversely, symmetric STDP with both LTP and LTD can be modeled by asserting that synaptic strengths are updated when either neuron spikes, with a change in synaptic strength from *l* to *k* given by the Mexican hat function (Ricker wavelet)

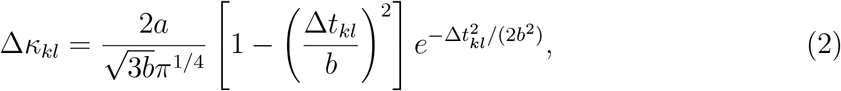

where *a* scales the magnitude of STDP, and *b* scales the temporal width of the Mexican hat (see Fig. 2A).

**Figure 2:**
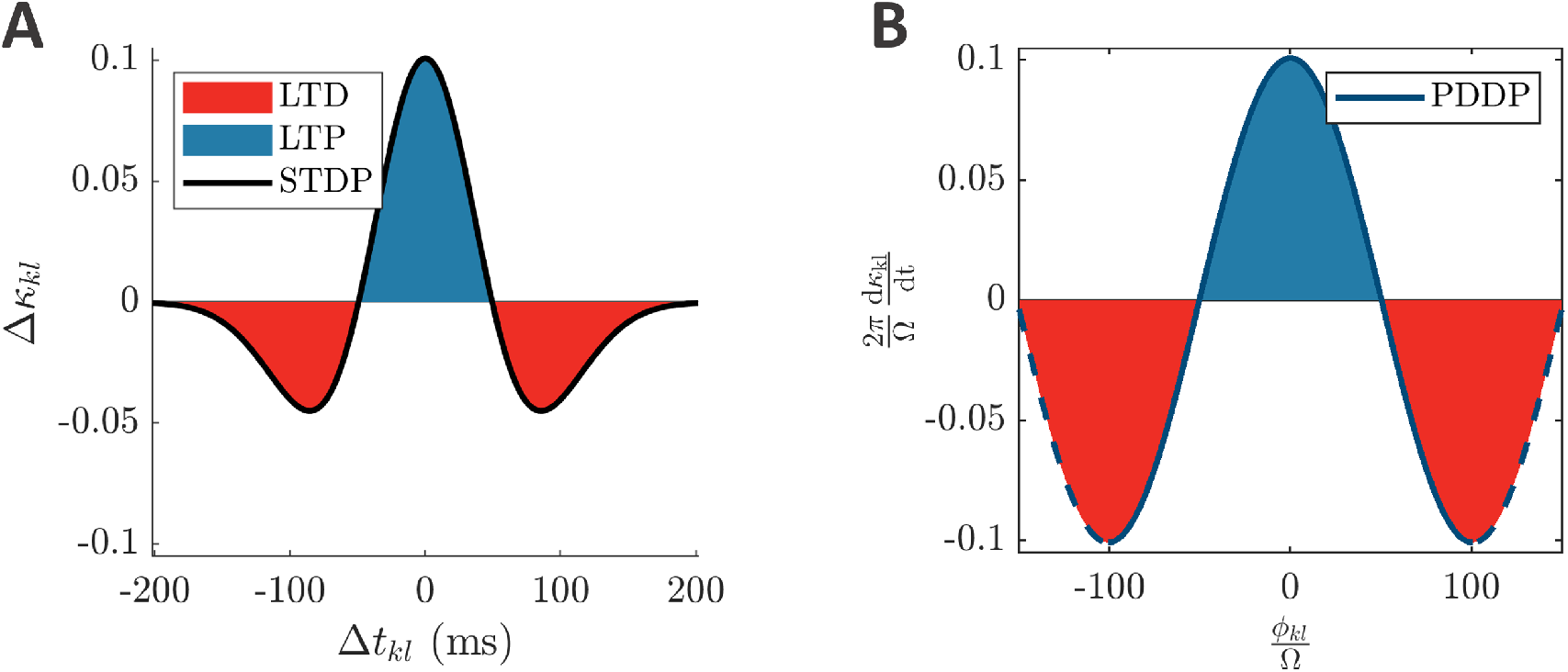
From symmetric STDP to single-harmonic PDDP. **A:** symmetric STDP function (Mexican hat, equation (2)) describing the change in weight *Δκ_kl_* as a function of the difference between spikes times Δ*t_kl_*. **B:** Considering the phase difference *ϕ_kl_* mod 2*π* instead of the difference between spike times, the STDP function in A can be approximated by the PDDP function in B (equation (3) with *φ* = 0). The horizontal axis is 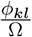, and the vertical axis is 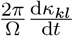 to enable comparison with panel A. The solid blue line correspond to one oscillatory period at frequency Ω centered on *ϕ_kl_* = 0, the dashed blue line extends beyond one period. The parameters used in panel A are *a* = 0.025822, *b* = 0.049415, corresponding to λ = 1, *∊* = 0.5, Ω = 10*π* in panel B. LTP is highlighted in blue, and LTD in red.

To approximate STDP in phase oscillator networks, simpler phase-dependent plasticity (PDDP) rules have been developed, which prescribe synaptic changes based on differences in phase of neuron pairs rather than differences in spike timing. If the state of oscillator *k* is given by a phase variable *θ_k_* ∈ [0, 2*π*) on the circle, the change of coupling weight between oscillators *k* and *l* according to PDDP depends on their phase difference *θ_l_* – *θ_k_*. While symmetric STDP can be approximated by the symmetric PDDP rule originally proposed by Seliger *et al.* (Seliger, Young, & Tsimring, 2002), several authors added a phase-shift parameter *φ* in order to approximate causal STDP-like learning (Aoki & Aoyagi, 2009; Berner et al., 2019), as well as other types of plasticity. With this single-harmonic PDDP rule, the weight *κ_kl_* from oscillator *l* to oscillator *k* evolves according to

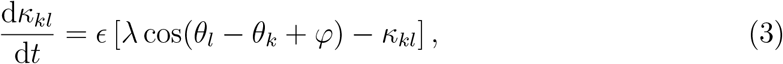

where λ controls the strength of PDDP relative to the decay of the synaptic strength and *ϵ* sets the relative time scale between the plasticity mechanism and the phase dynamics. When *φ* = 0, we recover the learning rule introduced by Seliger *et al.*, which is symmetric around a phase difference of zero. For *φ* = *π*/2, the cosine term is anti-symmetric around zero, which provides a first level of approximation of additive, causal STDP (in the absence of the decay term). However, this is a coarse approximation. A PDDP rule directly based on causal STDP exponential kernels (Maistrenko, Lysyansky, Hauptmann, Burylko, & Tass, 2007) could be more closely related to causal STDP. Lücken *et al.* proposed such a rule, and determined the correspondence between the parameters of the causal STDP rule, the parameters of the PDDP rule, and the parameters of the underlying network of coupled oscillators (Lücken, Popovych, Tass, & Yanchuk, 2016) (see Fig. 4B). Importantly, the synaptic strengths are continuously updated based on the evolving phase differences, while standard models of neural plasticity assume updates as discrete events when a neuron spikes.

Here, we construct mean-field approximations for coupled Kuramoto phase oscillators subject to PDDP and compare these approximations to fully adaptive networks where every edge evolves according to STDP. Specifically, we consider two types of PDDP rules: rules which update connection weights continuously as was done in previous studies, and event based rules, where weights are updated according to phase differences at a particular phase corresponding to spiking. In Section 2 we show that single-harmonic PDDP rules can indeed approximate symmetric STDP in adaptively coupled networks of Kuramoto oscillators, while a multi-harmonic rule is required to accurately approximate causal STDP. For PDDP rules, we derive exact equations describing the evolution of the mean coupling strength in terms of the Kuramoto/Daido order parameters that encode synchrony of the oscillators’ phases; see Section 3. We then focus on networks with symmetric adaptive coupling that naturally form clusters (cf. Section 2 or (Berner et al., 2019)). For such networks we construct mean-field approximations (Section 4) for the emergent coupling topologies, where each cluster corresponds to a coupled population. If we assume that coupling between clusters is through the mean coupling strength—rather than by individual weights between oscillators—we obtain low-dimensional Ott–Antonsen equations for the mean-field limit. We explicitly analyze the dynamics of the reduced equations for adaptive networks for one and two clusters. Note that these mean-field descriptions are not valid globally (i.e., there is no single reduced equation that is valid on all of state space) but rather aim to capture the dynamics on part of overall phase space determined by the initial conditions. In other words, we have a family of low-dimensional dynamics that can describe part of the phase space for this very high-dimensional system. Finally, we show that the dynamics of the full network can be approximated by such a family of low dimensional dynamics by extracting the mean field description from the emergent clustering (Section 5). Since brain activity is transient, we focus on transients and consider additive plasticity without bounds on individual weights for causal STDP. In line with previous studies, we however include a weight decay term when considering symmetric STDP (Seliger et al., 2002; Berner et al., 2019).

## 2 PDDP can approximate STDP in Kuramoto networks

Adaptation of network connections through STDP rely—as the name suggests—on the timing of action potentials of the coupled neurons. If the state of each neuron can be described by a single phase variable (for example, if the coupling is weak (Ashwin, Coombes, & Nicks, 2016)) then it may be possible to approximate STDP by an adaptation rule that depends on the phase differences between oscillators such as equation (3). In this section we now consider general PDDP rules, that can update weights continuously (as in equation (3)) or update at discrete time points (spiking events). We show that, for a network of phase oscillators, these PDDP rules can approximate both symmetric and causal STDP. For causal STPD, the accuracy increases substantially as the number of harmonics in the PDDP rule is increased.

To illustrate this, we focus on the Kuramoto model (Kuramoto, 1975), which is widely used to understand synchronization phenomena in neuroscience and beyond, subject to plasticity. Specifically, we consider *N* coupled Kuramoto oscillators where oscillators represent coupled neurons (Weerasinghe et al., 2019; Nguyen, Hayashi, Baptista, & Kondo, 2020; Weerasinghe, Duchet, Bick, & Bogacz, 2021). The phase *θ_k_* of oscillator *k* evolves according to

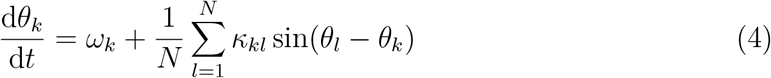

with intrinsic frequency *ω_k_* and strength *κ_kl_* of the synaptic connections from oscillator *l* to oscillator *k* (subject to plasticity). The (complex-valued) Kuramoto–Daido order parameters

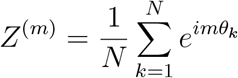

for 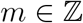 capture the (cluster) synchrony of the oscillator phases. The magnitude of the first order parameter *Z* ≔ *Z*^(1)^—simply called the Kuramoto order parameter—captures global synchrony, that is, for *Z* = *ρ*e^*i*Ψ^ we have |*Z*| = *ρ* = 1 if all oscillators have the same phase *θ*_1_ = · … = *θ_N_*. Similarly, |*Z*^(2)^| = 1 if the oscillators form two antiphase clusters where *θ_j_* = *θ_k_* or *θ_j_* = *θ_k_* + *π*, etc.

### 2.1 Principles to convert STDP to PDDP

PDDP rules prescribe synaptic changes based on differences in phase of neuron pairs rather than differences in spike timing, and can be used to approximate both symmetric and causal STDP in phase oscillator networks. In particular, the approximation is expected to hold under the assumption that the evolution of phase differences is slower than the phase dynamics (Lücken et al., 2016). Under this assumption, spike time differences are approximated by dividing phase differences by the mean angular frequency of the network 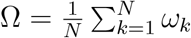.

As the phase difference is continuous in time, the discrete weight updates, based on spike-time differences in the case of STDP, can be converted to a continuous-time differential equation in terms of the phase differences. As a result, the coupling weight between each pair of neurons updates continuously based on the phase difference between the pre- and postsynaptic oscillators; we refer to this as continuously updating PDDP or simply PDDP when there is no ambiguity. STDP updates occur every time a neuron spikes, while PDDP updates occur continuously at every point in time. To ensure that STDP and continuously updating PDDP scale similarly, we scale the discrete STDP updates by the average number of spikes per unit time Ω*/*2*π*.

Rather than updating weights continuously, we can restrict weight updates to occur only at spiking events. We say that oscillator *k* spikes if its phase increases through *θ_k_* = 0, and let 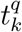 be the *q*^th^ firing time of neuron *k*. At each spiking event, we update the coupling weight between each pair of neurons based on their phase difference; we give explicit examples of the functional form of the updates below. We refer to this type of PDDP rule as event-based PDDP (ebPDDP). While there is an explicit phase dependence through the events, the actual change only depends on the phase difference. To ensure appropriate scaling, we again multiply by the average number of spikes per unit time and introduce an additional factor which is only non-zero when either the pre- or postsynaptic neuron spikes. This factor is defined as 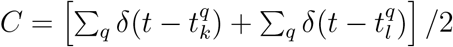, where *δ* denotes the Dirac delta function.^2^

### 2.2 Symmetric STDP and single-harmonic PDDP

In this section, we show that symmetric, non-causal STDP modelled by equation (2) together with the weight decay 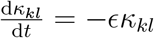 can be approximated by the single-harmonic PDDP learning rule introduced by Seliger *et al.* (equation (3) with *φ* = 0). We refer to this rule in what follows as the Seliger rule. As detailed in the previous section, discrete STDP updates should be scaled by *Ω*/2*π* to obtain continuous PDDP updates. Therefore, by matching the scaled maximum of equation (2) with the maximum of equation (3), we have 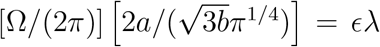, which determines the value of λ for a given *ϵ* (see example in Fig. 2). Moreover, to ensure that the scale of spike timing differences in the STDP rule and the scale of phases differences in the PDDP rule match without modifying the Seliger rule, we choose *b* ≈ *π*/(2Ω) (see Fig. 2). Arbitrary values of *b* could be accommodated by scaling the phase difference term in the Seliger rule as detailed in the previous section. The corresponding event-based PDDP rule reads

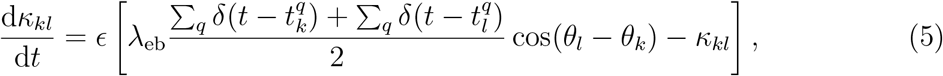

with 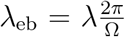. Note that updates to the coupling strengths happen whenever the pre- or postsynaptic neurons spikes.

As shown in Fig. 3, both single-harmonic PDDP rules (the Seliger rule and equation (5)) can approximate symmetric STDP (equation (2)) in networks of Kuramoto oscillators. Simulation details can be found in Section A.1. The time evolution of the weight distribution (panel A), the coupling matrix at the last simulation time point (panel B), as well as the the time evolution of the average coupling (panel C1) and network synchrony (panels C2-C3) are comparable across learning rules. In particular, the Pearson’s correlation between the STDP coupling matrix and the PDDP coupling matrix is 0.88, while the correlation between the STDP coupling matrix and the ebPDDP coupling matrix is 0.90. For the parameter set shown in Fig. 3, we see the emergence of two synchronised clusters. For this type of dynamics, the average coupling as a function of time may be better captured by ebPDDP than PDDP (panel C1). However, for the desynchronized state (shown in the Supplemental Material C Fig. C.1), there is little difference between continuous PDDP and ebPDDP. In both states, the accuracy of the approximation could be improved by considering a Fourier expansion of the Mexican hat function rather than a single cosine term. We explore this for causal, non symmetric STDP in the next section, as we found that a single sine term is, in general, a poor approximation of the causal STDP kernel.

**Figure 3:**
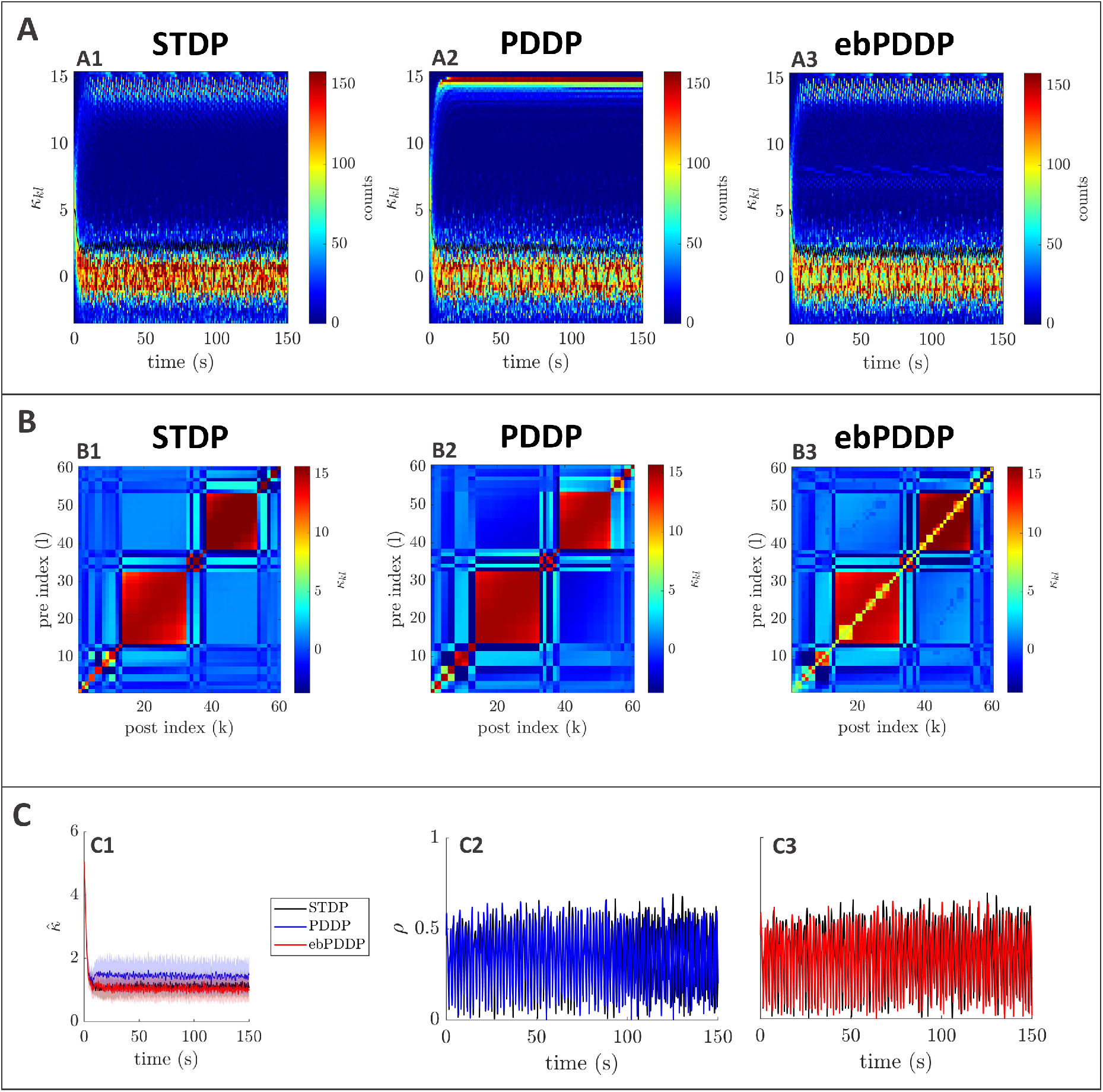
Comparison between symmetric STDP and PDDP in a Kuramoto network (two synchronised cluster state). **A:** Evolution of the distribution of coupling weights with time (100 bins at each time point). The average weight is represented by a thin black line. STDP is shown in A1, PDDP in A2, and ebPDDP in A3. **B:** Coupling matrix at *t* = 150s, with oscillators sorted by natural frequency. STDP is shown in B1, PDDP in B2, and ebPDDP in B3. **C:** Time evolution of average coupling (C1, error bars represent the standard error of the mean over 5 repeats, the high variability is due to sensitivity to initial conditions) and network synchrony (C2-C3). STDP is shown in black, PDDP in blue, and ebPDDP in red. [*a* = 0.38733, *b* = 0.049415, *σ_κ_* = 3, Δ = 1.2*π*, Ω = 10*π* (5Hz)]

### 2.3 Causal STDP and multi-harmonic PDDP

In the previous section, we considered PDDP rules with a single harmonic in the phase difference. To get a better approximation of causal STDP, one can take more harmonics into account.

#### 2.3.1 Obtaining multi-harmonic PDDP rules from causal STDP

To approximate causal STDP, we consider the PDDP rule proposed by Lücken *et al.* (Lücken et al., 2016) as

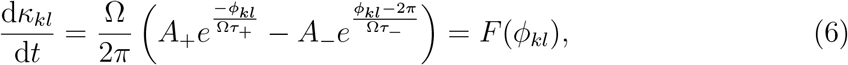

where *ϕ_kl_* = (*θ_l_*–*θ_k_*) mod 2*π*, Ω is the mean (angular) frequency of the network, and other parameters have been defined in equation (1). In equation (6), both synaptic potentiation (first term) and synaptic depression (second term) are described without requiring a piecewise definition thanks to the fast decaying exponentials. The correspondence of this PDDP rule with additive STDP is illustrated in Fig. 4B. If the postsynaptic neuron *k* spikes (i.e., its phase increases through 0) shortly after the presynaptic neuron *l*, *ϕ_kl_* is small and positive, which will lead to a potentiation of *κ_kl_*. Conversely, if the postsynaptic neuron spikes shortly before the presynaptic neuron, *ϕ_kl_* – 2*π* is small and negative, which will lead to a depression of *κ_kl_*. As laid out in Section 2.1, spike time differences are approximated in equation (6) by dividing phase differences by the network mean angular frequency. Moreover, the scaling factor Ω/2*π* (average number of spikes per unit time) accounts for the conversion of discrete weight updates to a continuous-time differential equation. The corresponding event-based PDDP rule can be obtained as

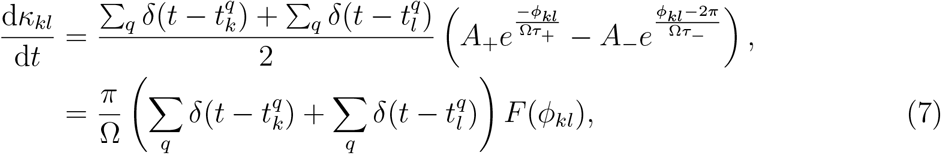

where *F* is given by equation (6).

**Figure 4:**
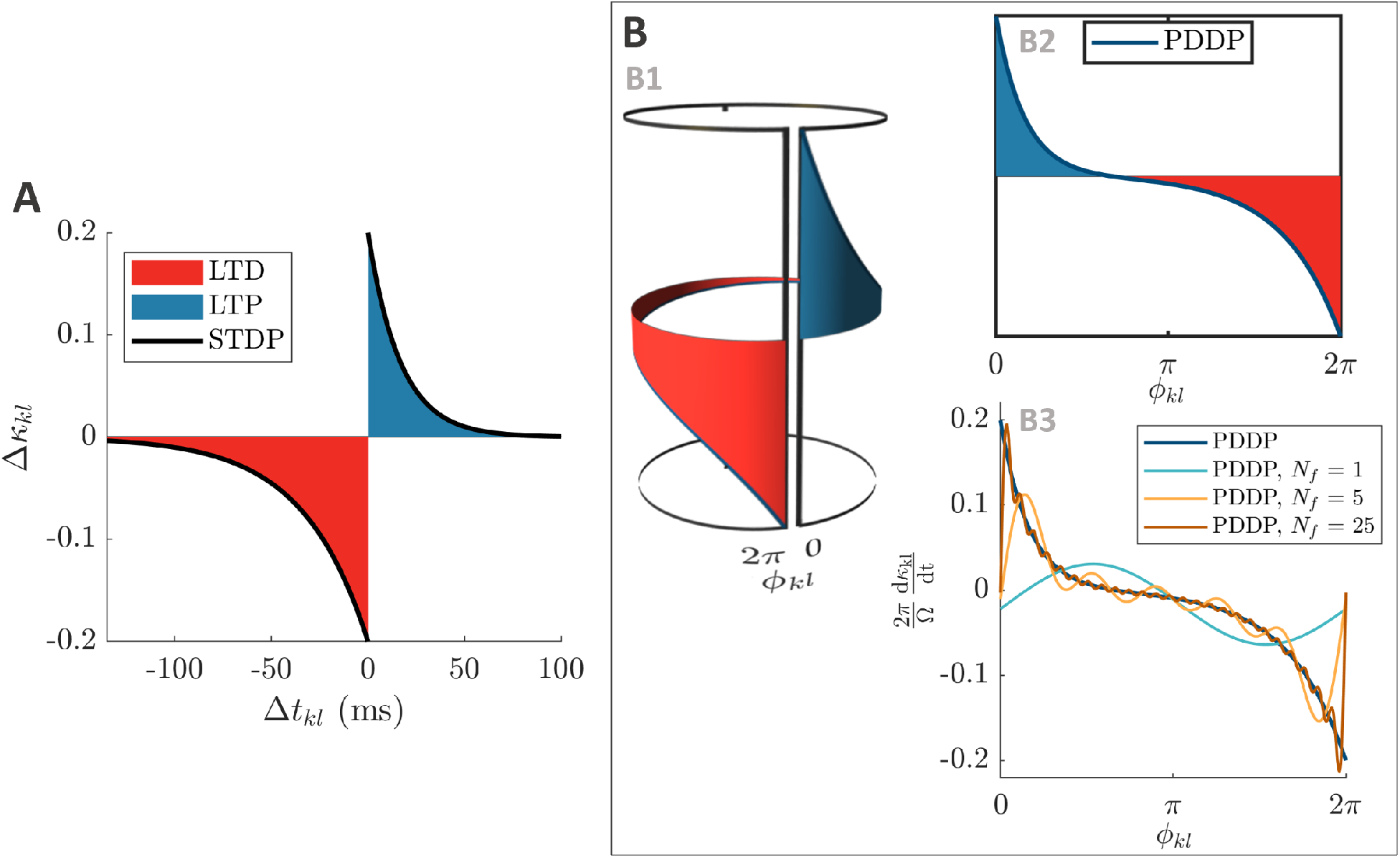
From causal STDP to multi-harmonic PDDP. **A:** STDP function (equation (1)) describing the change in weight Δ*κ_kl_* as a function of the difference between spikes times Δ*t_kl_*. **B:** Considering the phase difference *ϕ_kl_* mod 2*π* instead of the difference between spike times, the STDP function in A can be approximated by the PDDP function in B2 (equation (6)). As shown in B1, wrapping the PDDP function around a cylinder to join *ϕ_kl_* = 0 and *ϕ_kl_* = 2*π* illustrates the correspondence with the STDP function. In B3, the PDDP function is approximated using truncated Fourier series with 1, 5, and 25 Fourier components. The vertical axis is 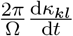 to enable comparison with panel A. The parameters used in all panels are *τ*_+_ = 16.8ms and *τ*_ − = 33.7ms, *A*_+_ = *A*___ = 0.2. LTP is highlighted in blue, and LTD in red.

In both cases, we can expand *F*(*ϕ_kl_*) as a Fourier series of the phase difference since *ϕ_kl_* is defined modulo 2*π*, which makes *F*(*ϕ_kl_*) a 2*π*-periodic function. The Fourier expansion will be key to derive the evolution of the average coupling strength in Section 3, and can be truncated to only include *N_f_* components for simulations (see examples in Fig. 4C). We note that the PDDP rule by Berner *et al.* with *φ* ≈ *π*/2 is a single-harmonic version of the rule by Lücken *et al.* (with a vertical shift), and call truncated Fourier expansions of equation (6) and equation (7) with *N_f_* > 1 “multi-harmonic PDDP”. We express the Fourier series of *F* as

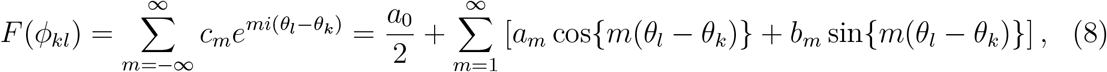

where 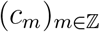 are the complex-valued Fourier coefficients, or equivalently 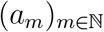 and 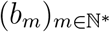 are the real-valued Fourier coefficients. The Fourier coefficients only depend on the parameters of the STDP rule (equation (1)) and Ω, and the real-valued coefficients can be obtained analytically as

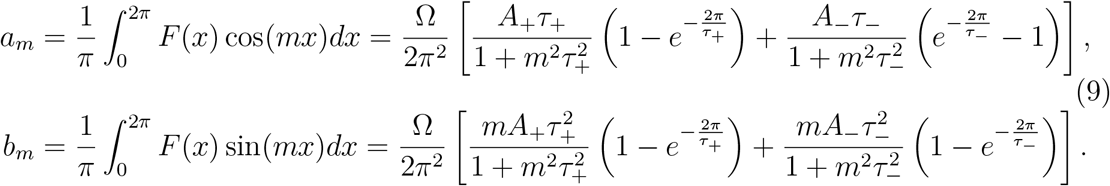

#### 2.3.2 Comparison of learning rules

Multi-harmonic PDDP can approximate causal STDP in Kuramoto networks (simulation details can be found in the Supplemental Material A.2). As the number of harmonics included in the PDDP rules is increased, the dynamics for the networks with PDDP begin to match those of the STDP network (Fig. 5). The time evolution of the weight distribution (panel A), the coupling matrix at the last simulation time point (panel B), as well as the the time evolution of the average coupling (panel C1) and network synchrony (panels C2-C7) are closely matched for causal STDP and multi-harmonic PDDP or ebPDDP when enough Fourier components are included (*N_f_* = 25). Simulations were also performed for a range of parameter values and the findings were similar (see Figs. C.2–C.5 in Supplemental Material C). Single-harmonic PDDP or ebPDDP (*N_f_* = 1) can provide a first level of approximation of causal STDP in certain cases when the time evolution of the weight distribution is simple (see Supplemental Material C Fig. C.4). However, single-harmonic rules are unable to describe more complex cases, even qualitatively (as seen in Fig. 5). In all cases studied for a network frequency of 5Hz, 25 Fourier components are deemed sufficient to approximate the dynamics of the network with causal STDP. While the performance of ebPDDP is similar to PDDP, ebPDDP is slightly more accurate than PDDP. To quantitatively compare STDP to PDDP and ebPDDP, we construct error metrics for the time evolution of the weight distribution *e*_hist_(_*κkl*_), the average coupling 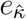, and the network synchrony *e_ρ_*, and consider the Pearson’s correlation between coupling matrices at the last simulation time point 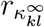. These metrics are defined in the Supplemental Material A.2. In general, these metrics improve with increasing *N_f_* (Fig. 6A), although stagnation or a slight worsening can be seen when the error is already low. Although it is of no consequence in Kuramoto networks with sine coupling, self-coupling weights are consistently different between causal STDP and multi-harmonic PDDP or ebPDDP in our simulations. This is due to the truncated Fourier expansions of *F* not being zero when the phase difference is zero (see Fig. 4C).

**Figure 5:**
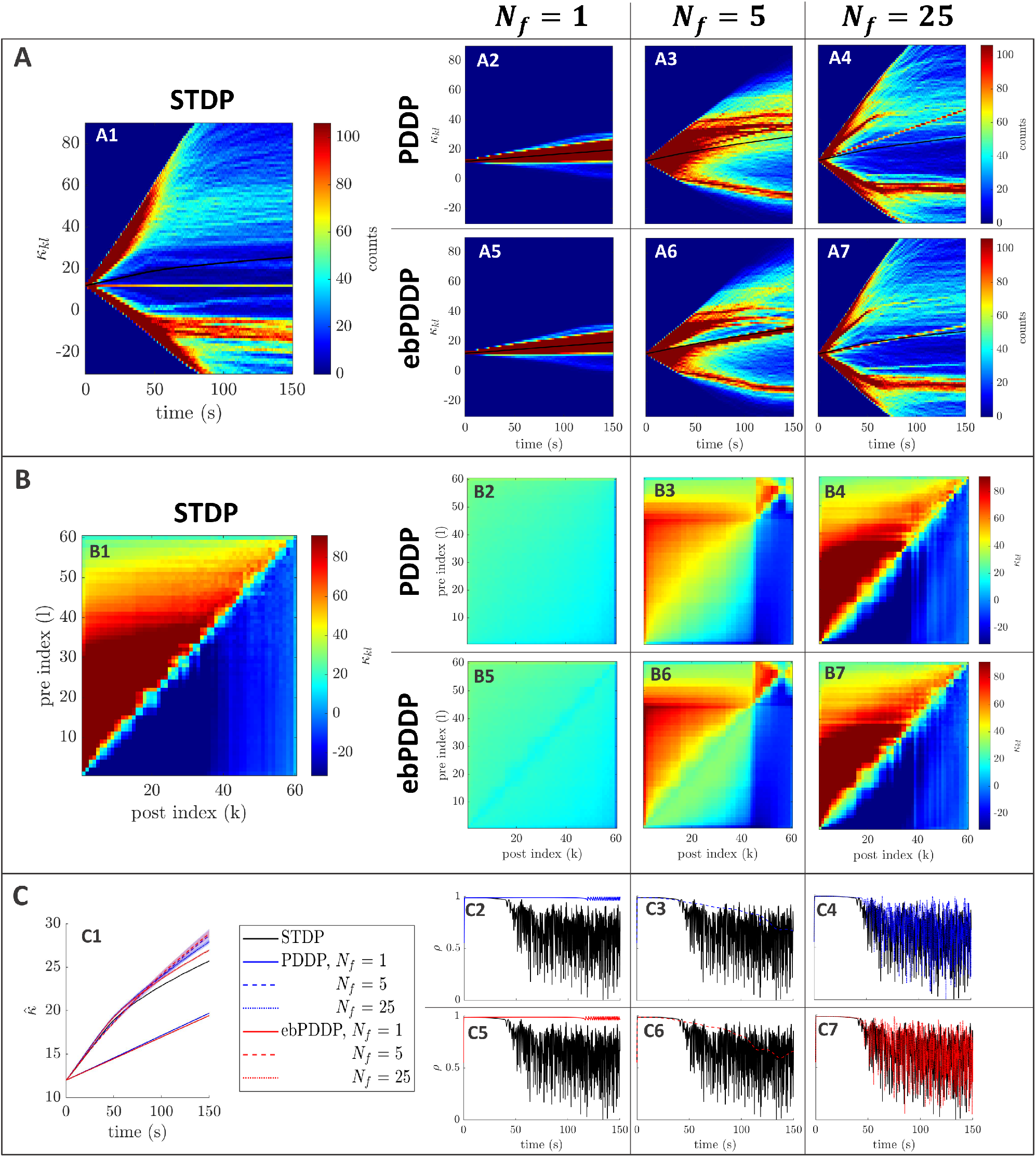
Comparison between STDP and PDDP in a Kuramoto network, [*β* = 0.5, *σ_κ_* = 0.2, Δ = 0.6*π*, Ω = 10*π* (5Hz)]. Results for PDDP and ebPDDP are shown for 1, 5, and 25 Fourier components *Nf* (first, second, and third column on the right hand side of the figure, respectively). **A:** Evolution of the distribution of coupling weights with time (100 bins at each time point). The average weight is represented by a thin black line. STDP is shown in A1, PDDP in A2-A4, and ebPDDP in A5-A7. **B:** Coupling matrix at *t* = 150s, with oscillators sorted by natural frequency. STDP is shown in B1, PDDP in B2-B4, and ebPDDP in B5-B7. **C:** Time evolution of average coupling (C1, error bars represent the standard error of the mean over 5 repeats) and network synchrony (C2-C7). STDP is shown in black, PDDP in blue, and ebPDDP in red. In all panels, the approximation becomes better as *N_f_* is increased.

**Figure 6:**
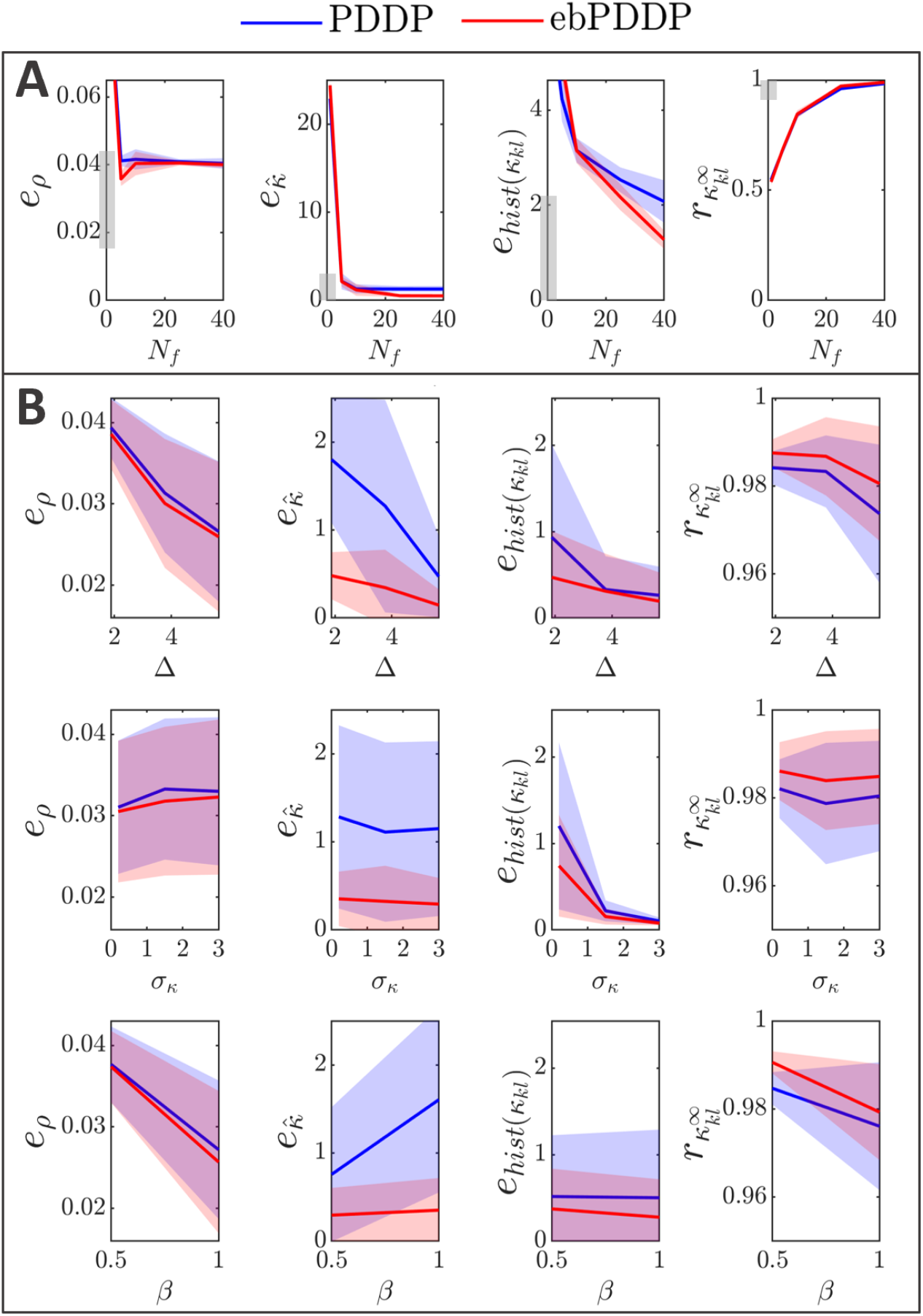
Influence of *N_f_*, *σ_n_*, Δ, and *β* on error metrics. Error metrics for PDDP compared to STDP for the time evolution of network synchrony *e_p_*, average coupling 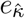 and distribution of weights *e*_hist(*κ_kl_*)_, and for the coupling matrix at the last stimulation point 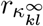 are shown in the first, second, third, and fourth columns, respectively. Results for PDDP are in blue, and for ebPDDP in red. **A:** Influence of the number of Fourier coefficients *N_f_* on the error metrics for the parameters used in Fig. 5 (*β* = 0.5, *σ_κ_* = 0.2, Δ = 0.6*π*, Ω= 10*π*), error bars show the standard error of the mean (sem) over 5 repeats. **B:** Influence of the network parameters on the error metrics for *N_f_* = 40. The standard deviation of the oscillator frequency distribution Δ is shown in the first row, the standard deviation of the initial weight distribution *σ_κ_* is given in the second row, and the ratio the LTP to LTD scaling factors *β* is depicted in the third row. All of the combination of parameters Δ = {0.6*π*, 1.2*π*, 1.8*π*}, *σ_κ_* = {0.2, 1.5,3}, *β* = {0.5, 1} are included with 5 repeats for each combination. In each row, averaging is performed over the parameters that do not correspond to the horizontal axis (standard deviation error bars). Note that the scale of the vertical axes is two to ten times smaller than in panel A for readability (range indicated by grey bars). See Supplemental Material C Fig. C.9 for detailed slices in parameter space.

#### 2.3.3 Parameter dependence

Model parameters influence the four metrics described in the previous section (Fig. 6B), although the impact on 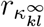 is minor (always stays > 0.96). Increasing the standard deviation of the frequency distribution (Δ) tends to improve the error metrics, except 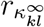 which gets slightly lower. This is expected since a larger Δ reduces synchrony and makes the weight distribution more unimodal. The impact of the standard deviation of the initial coupling distribution (*σ_κ_*) on the metrics is smaller, except for the time evolution of the weight distribution. The effect of the ratio of the scales of LTD to LTP (*β* = *A*_ − /*A*_+_)depends on the metric considered. The largest effect is a lowering of *e_ρ_* when LTD dominates (*β* = 1) compared to the balanced situation (*β* = 0.5). This is due to the fact that the time evolution of *ρ* is closely approximated with *N_f_* = 1 when LTD dominates, whereas in the balanced state more Fourier components are required to obtain a good approximation. Since dominant LTD leads to lower synchrony, this matches the previous observation that lower synchrony is associated with lower error metrics. Time courses of *ρ* for both states can be found in Supplemental Material C; Fig. C.2 shows the balanced state and Fig. C.4 shows the LTD dominant regime.

To test the robustness of our findings, we studied a network with a mean frequency four times higher (20Hz) and considered the least favorable part of parameter space (lowest Δ, lowest *σ_κ_*, and *β* = 0.5) (see Supplemental Material C Fig. C.5). The order of magnitude of the error metrics is the same as for 5Hz, except for 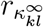 (Supplemental Material C Fig. C.8C). The lower value for 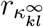 may be explained by the greater complexity and finer structures in the coupling matrix. However, the coupling matrix obtained at the last stimulation point with multi-harmonic ebPDDP for *N_f_* = 75 (see Supplemental Material C Fig. C.5B7) is qualitatively very similar to the coupling matrix obtained with STDP (panel B1). At 20Hz, the period of the oscillators is comparable to the STDP time constants, hence the weight distribution patterns unfolding in time are more highly multimodal than at 5Hz. These patterns are still well approximated by multiharmonic PDDP and ebPDDP, but a larger number of Fourier components than at 5Hz is warranted for accurate results. Simulating this higher frequency network required a smaller simulation time step (Δ*t* = 0.1ms). However, the time step has overall little impact on the error metrics at lower frequencies (Fig. C.8B).

Additional explorations of the parameter dependencies can be found in the Supplemental Material C Fig. C.8 and Fig. C.9.

## 3 Evolution of the average coupling strength in networks with PDDP

While each coupling weight *κ_kl_* evolves independently of one another, we now derive evolution equations for the average coupling weight

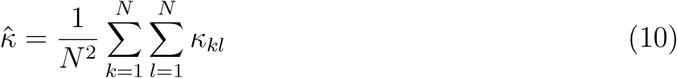

for both continuously updating and event-based PDDP. Obtaining the mean coupling weight dynamics is necessary for constructing our mean-field approximations, which are based on the average coupling weights within and between populations.

### 3.1 Average coupling for general PDDP

Suppose that each weight *κ_kl_* evolves according to a general, continuously updating PDDP rule 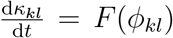, where *F* is a 2*π*-periodic function of the phase difference *ϕ_kl_*. The PDDP rule in Section 2.3 with *F* approximating causal STDP (equation 6) is a particular example. Writing 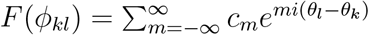 as in (8) and differentiating (10) yields

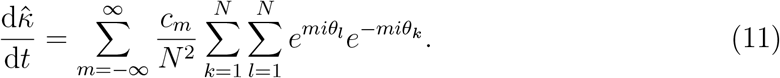

This expression can be written in terms of the Kuramoto–Daido order parameters *Z*^(*m*)^. We have 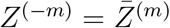 and consequently Eq. (11) reads

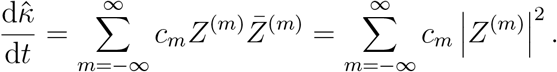

The series converges as |*Z*^(*m*)^| ≤ 1 and the Fourier series of *F* is assumed to converge. Since *c_m_* + *c*__*m*_ = *a_m_* and 2*c*_0_ = *a*_0_, the evolution equation for 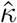 can be simplified to

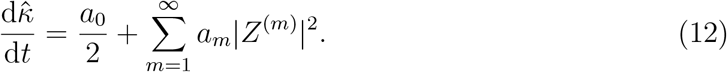

In the case of *F* approximating causal STDP, the coefficients 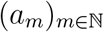 are given by equation (9). In the absence of bounds on the coupling weights, equation (12) exactly describes the average coupling strength in a phase oscillator networks with PDDP as illustrated in Fig. 7.

**Figure 7:**
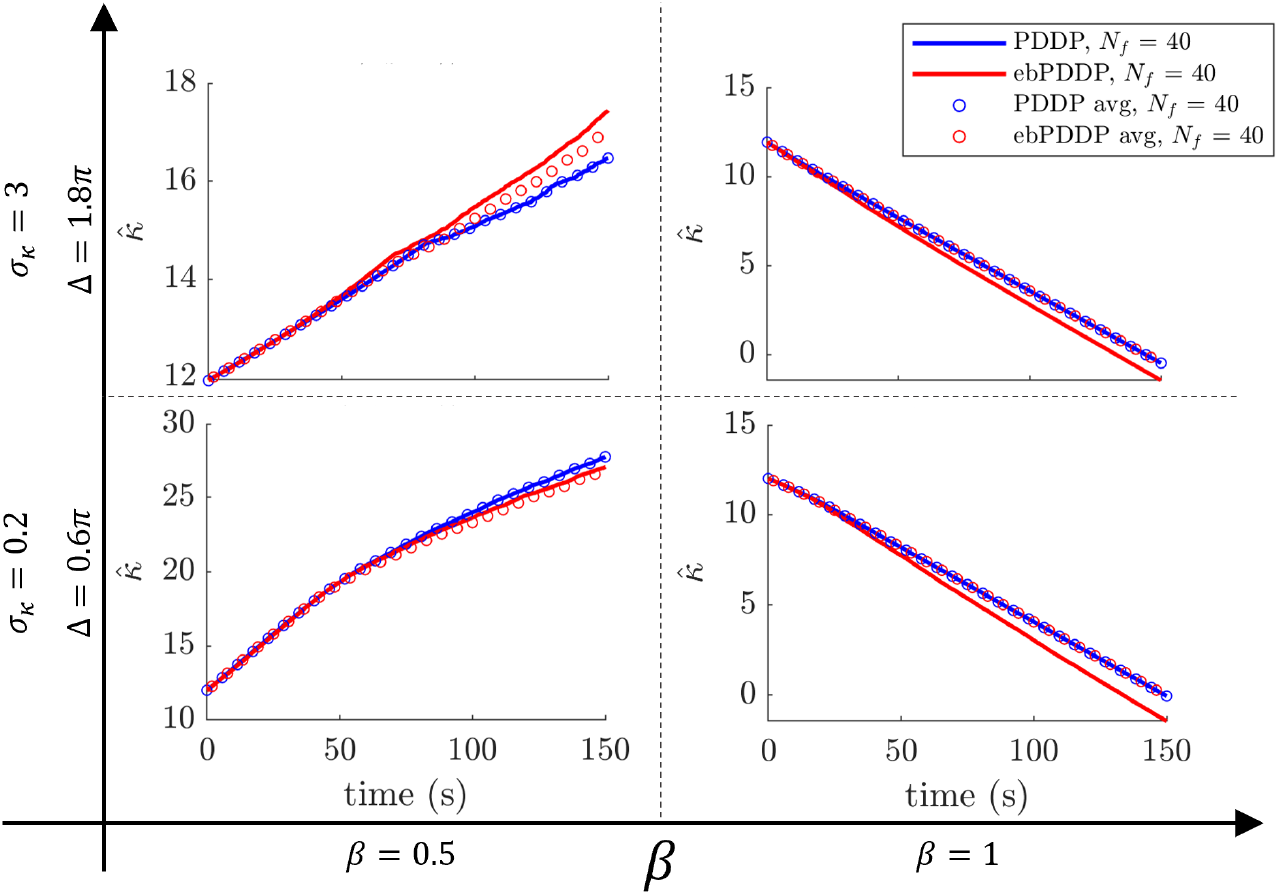
Simulation of average coupling rules in Kuramoto networks. Equation (12) (blue circles) describes exactly the average coupling weight in Kuramoto networks with PDDP (blue lines). The correspondence between equation (15) (red circles) and the average coupling weight in Kuramoto networks with ebPDDP (equation 7, red lines) is not exact as explained in the main text. The same number of Fourier components are used in all cases (*N_f_* = 40). The four sets of parameters used for the simulations are indicated by the bold axes and correspond to those used in Fig. 5 as well as Fig. C.2, C.3, and C.4 from the Supplemental Material C.

If the PDDP rule contains a decay term, the evolution of the mean coupling strength will reflect this as well. More concretely, consider an evolution of individual coupling weights 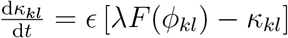 as in the PDDP rule by Seliger *et al.* (Seliger et al., 2002) where λ and *ϵ* are parameters. Then the evolution of the average coupling is

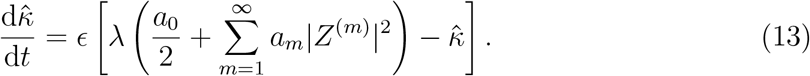

In particular, for *a*_1_ = cos(*φ*) and all other Fourier coefficients equal to zero, this equation exactly describes the evolution of the average coupling for the PDDP rule given by equation (3).

### 3.2 Average coupling for general event-based PDDP

We consider the general ebPDDP rule represented by equation (7) where *F* is any 2*π*-periodic function of the phase difference that can be expanded as a Fourier series according to equation (8). To obtain the corresponding average coupling strength, the Dirac deltas indicating spiking events need to be expressed as functions of neuron’s phases. Since *θ_k_* is defined mod 2*π*, we have 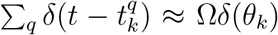 as in (Coombes & Byrne, 2019). We use this approximation to define an ebPDDP rule which depends only on the phases of the oscillators,

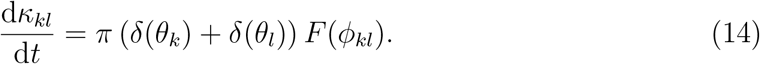

Using the Fourier expansion of *F* as before, the corresponding average coupling 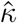 can be obtained as

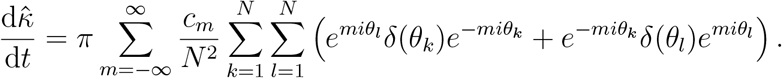

With *θ* defined mod 2*π*, *δ*(*θ*) can also be Fourier-expanded as 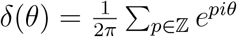 (Coombes & Byrne, 2019). Since

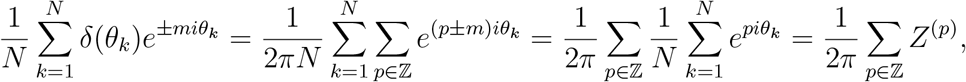

we obtain

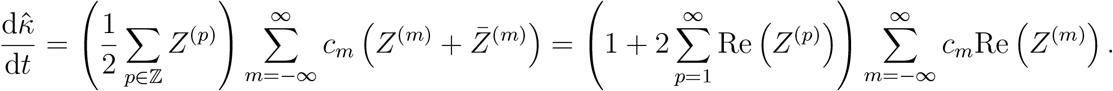

Using the real-valued Fourier coefficients defined in equation (9), this expression becomes

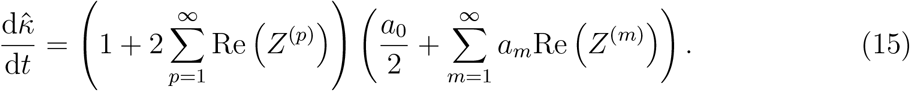

As in the previous subsection, this result can be extended to include a decay term.

Simulations show that average weights obtained from equation (15) are similar to average weights obtained from equation (7) as shown in Fig. 7. Although equation (15) is an exact description of the average weight in phase oscillator networks with adaptivity given by equation (14) (where the Dirac deltas are functions of phase), our simulations are based on equation (7) (where the Dirac deltas are functions of time). The approximation 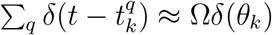 used to derive equation (14) from equation (7) gives rise to the discrepancies visible in Fig. 7.

## 4 Mean-field dynamics of oscillator populations with adaptive coupling

Symmetric or nearly symmetric STDP rules can lead to the formation of densely connected clusters with homogeneous coupling strength; cf. Section 2.2 and, e.g., (Popovych,Xenakis, & Tass, 2015; Berner et al., 2019; Röhr, Berner, Lameu, Popovych, & Yanchuk, 2019). Specifically, Figure 3 shows the emergence of multiple clusters of distinct mean intrinsic frequencies. For the remainder we will focus on such plasticity rules and interpret each cluster as an emergent population of phase oscillators. Suppose that there are *M* emergent clusters and the corresponding populations *μ* ∈ {1,…, *M*} have *N_μ_* oscillators. Rewriting (4) with the cluster labelling, the evolution of *θ_μ,k_*, the phase of oscillator *k* ∈{ 1, …, *N_μ_*} in cluster *μ*, evolves according to

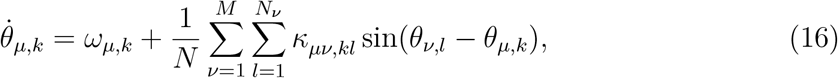

where *κ_μν,kl_* is the coupling strength from oscillator *l* in population *ν* to oscillator *k* in population *μ*. We now suggest a low-dimensional description of the resulting dynamics for populations corresponding to emergent clusters in the fully adaptive network in terms of the population Kuramoto order parameters 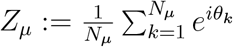.

### 4.1 Low-dimensional dynamics for homogeneous coupling

Using the assumption that the emergent coupling within and between clusters is homogeneous, we replace individual coupling strengths *κ_μν,kl_* from oscillators in population *ν* to oscillators in population *μ* by the mean coupling strength from population *ν* to population *μ*,

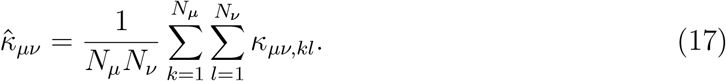

Writing 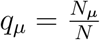 for the relative population size, we obtain

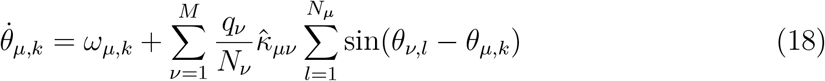

that describes homogeneously coupled populations.

Such networks of Kuramoto oscillators admit an exact low-dimensional description in terms of the dynamics of the population order parameter *Z_μ_* due to the Ott–Antonsen reduction (Ott & Antonsen, 2008, 2009); see also the recent review (Bick et al., 2020). In the mean-field limit of infinitely large networks, the Kuramoto–Daido order parameters 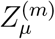 for each population *μ* describe the distribution of oscillators. The key observation for this reduction is that for networks of the form (18) the *m*^th^ Kuramoto–Daido order parameter can be expressed as a power of the Kuramoto order parameter 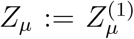, that is, 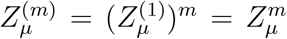. If we assume that the intrinsic frequencies *ω_μ,k_* are distributed according to a Lorentzian with mean Ω*_μ_* and width Δ_*μ*_, then the dynamics of (18) are determined by

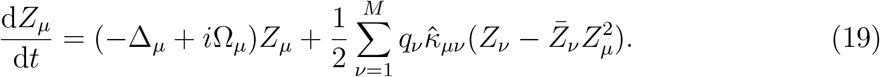

The dynamical equations for the evolving coupling weights can be derived as in Section 3 and then taking the limit *N* → ∞. Now the Ott–Antonsen reduction allows us to simplify the expressions through 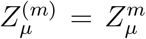. If individual weights evolve according to the general PDDP rule 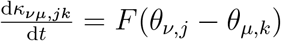 then

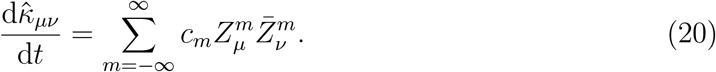

Similarly, for the event-based rule (14) we note that 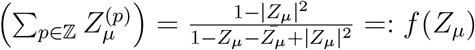 and thus

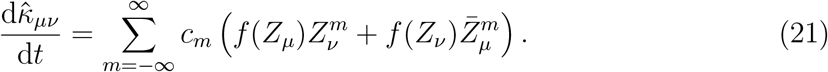

Decay terms can be incorporated in the same way as above.

Note that (19) together with either (20) or (21) form a closed set of equations. In the following we will analyze the dynamics of the reduced equations explicitly. Here, we will focus on the symmetric STDP rule approximated by a single harmonic PPDP rule (3) with *φ* = 0 such that

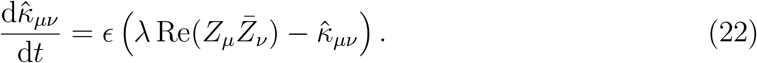

Equations (19) and (22) now form a closed low-dimensional system of coupled adaptive oscillator populations.

### 4.2 Single harmonic PDDP, one population

We begin by considering a one-population model to illustrate the types of behaviour possible for a one-cluster state. For a one-population model, the learning rule (22) does not depend on the mean phase. Thus, the phase dynamics decouple leading to two-dimensional effective dynamics^3^ determined by

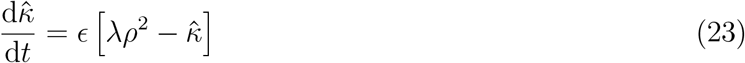

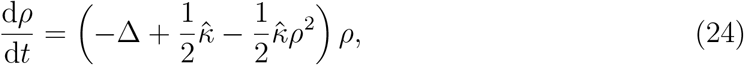

where *ρ* ≔ |*Z*|. Note that, in contrast to mean-field descriptions of Kuramoto oscillators with an adaptive global coupling parameter that depend linearly on the mean field *Z* (Ciszak, Marino, Torcini, & Olmi, 2020), we here have a quadratic dependency.

There is a trivial fixed point at 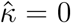, *ρ* = 0. While 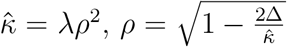 defines a pair of non-trivial fixed points, which exist for λ > 8Δ. Computing the Jacobian, we find that the trivial solution is stable for all physical parameter values (Δ > 0, *ε* > 0), and for the non-trivial fixed points, one is stable and the other unstable.

Using XPPAUT (Ermentrout, 2002), we performed a one parameter continuation in the heterogeneity parameter Δ (Fig. 8). When the heterogeneity is low, there exists a non-trivial stable fixed point, where the mean coupling does not decay to zero. Whereas after the saddle-node bifurcation at Δ = 0.125, the mean coupling will always decay to zero and the oscillators will be asynchronous (*ρ* = 0). As the strength of the plasticity rule λ is increased, the saddle-node moves to the right and the region of bistability (where the trivial and non-trivial fixed points co-exist) increases. These results agree with analytical results outlined above.

**Figure 8:**
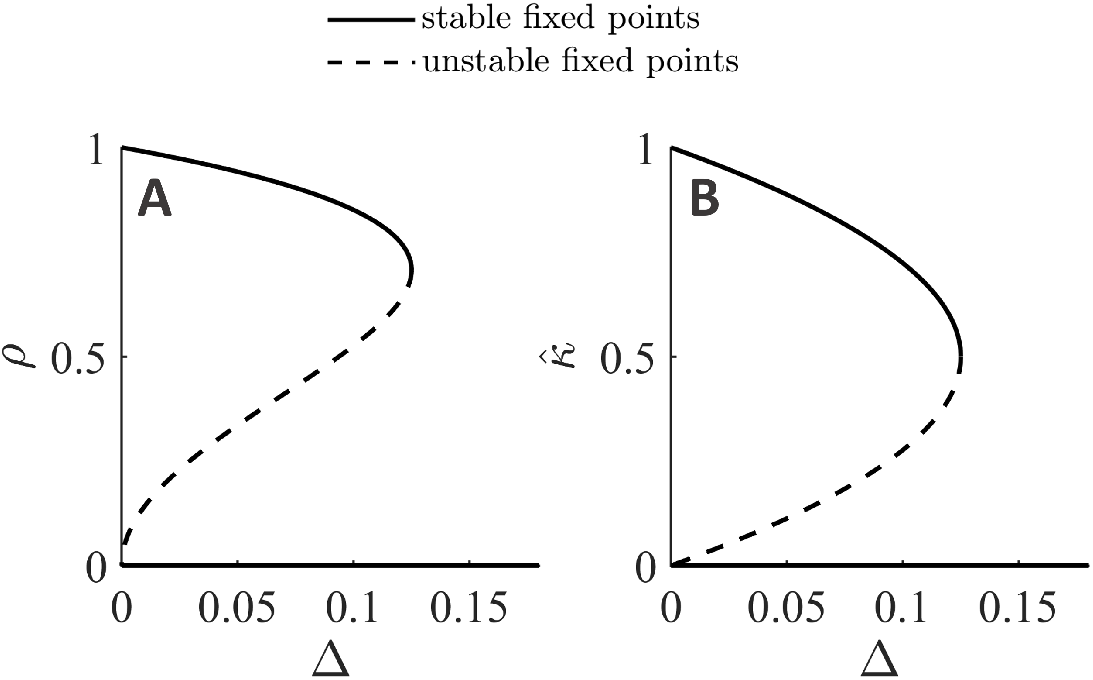
Bifurcation analysis of the mean-field equations for a single population. One parameter continuation in the heterogeneity parameter Δ for (23)–(24) with *λ* = 1 and *∈* = 0.5. A saddle-node bifurcation occurs at Δ = 0.125, which corresponds to the analytical value λ/8.

### 4.3 Single harmonic PDDP, two populations

Next consider two adaptively coupled populations evolving according to

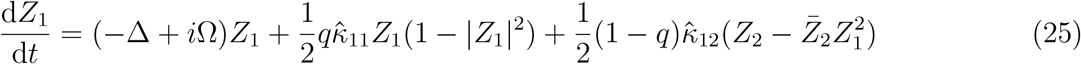

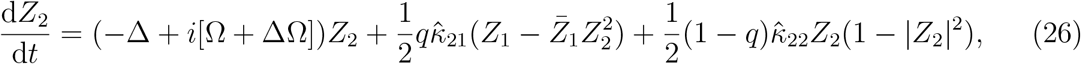

where *q* is the fraction of oscillators in population 1 and ΔΩ is the difference in mean intrinsic frequency between oscillators in each population. The dynamics for the intrapopulation coupling are as in the one population model,

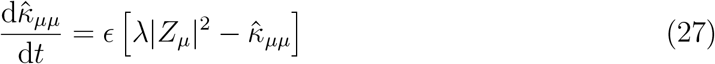

for *μ* ∈ [1, 2], while the inter-population coupling is given by (22).

Setting *q* = 0.5 (two equally sized populations), we perform a one parameter continuation in the intrinsic frequency difference ΔΩ (Fig. 9A). We find that for a small range of ΔΩ values (ΔΩ ∈ [−0.23, 0.23]) the two populations synchronize in frequency. This *frequency-locked solution* lies on an invariant set, where the level of synchrony of each population is identical (*ρ*_1_ = *ρ*_2_) and the coupling strengths are symmetric 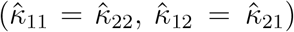. The system itself, however, is generally not symmetric (unless ΔΩ = 0) due to distinct intrinsic frequencies resulting in distinct mean phases of the two populations. The two non-trivial equilibrium branches in the inter-population coupling strength (Fig. 9A3), correspond to the in-phase/symmetric solution (positive 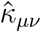) and the anti-phase/asymmetric solution (negative 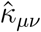).

**Figure 9:**
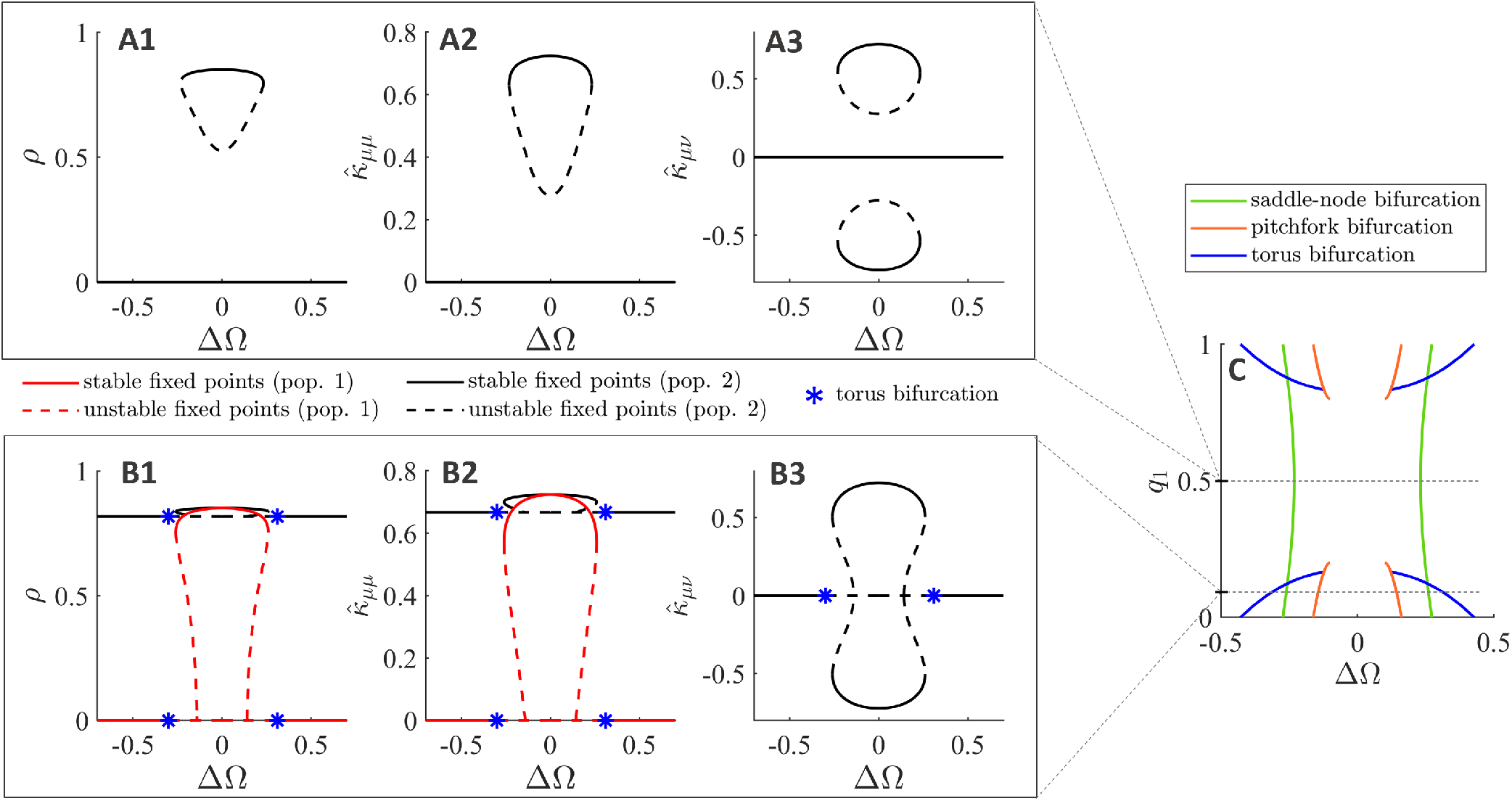
Bifurcation analysis of the mean field equations for a two population model. One and two parameter continuations for the system of equations given by (25) – (27) and (22). **A:** One parameter continuation in intrinsic frequency difference ΔΩ for equally sized populations (*q* = 0.5). Given the symmetry in the system, both populations have the same within-population synchrony and intra-/inter-population coupling strengths. Black solid (dashed) lines correspond to the stable (unstable) fixed point values for both populations/sets of coupling strengths. **B:** Continuation in ΔΩ for *q* = 0.1. As the mean inter-population coupling strengths are equal, the curve in panel B3 corresponds to both 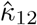 and 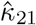. **C:** Two parameter continuation in the intrinsic frequency difference ΔΩ and the relative size of population 1 *q*, showing the saddle-node, pitchfork and torus bifurcation curves. Parameter values: Δ = 0.1, Ω = 30, *∈* = 0.5, λ = 1.

For general population sizes *q*, an additional branch of solution can emerge, where one population is fully asynchronous and the other is partially synchronized with non-trivial dynamics (e.g., *ρ*_1_ = 0, *ρ*_2_ ≠ 0); we call this the *decoupled solution*. More specifically, the structure of the equations implies that 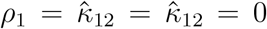 defines a dynamically invariant set. On this set, the coupling strength 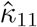 decays exponentially and the second population evolves independently of the first one (as the network is decoupled) according to the equations of motion for a single population with adaptive coupling. The decoupled solution now corresponds to the non-trivial solution branch for a single population (Fig. 8), which exists for λ > 8Δ. The only difference here is that the coupling 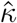 is replaced by an effective coupling 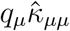 scaled with the (relative) population size. Hence, decoupled solutions only exists for *q* < 2Δ/λ (population 2 has non-trivial dynamics *ρ*_2_ ≠ 0) and *q* > 8Δ/λ (population 1 has non-trivial dynamics *ρ*_1_ ≠ 0). Note that these are equilibrium points for the effective dynamics but periodic solutions in the full system.

Letting *q* = 0.1, we find and continue the decoupled solution where population 1 is asynchronous and population 2 has non-trivial dynamics (*ρ*_1_ = 0, *ρ*_2_ ≠ 0) (Fig. 9B). The dynamics of population 1 (population 2) is shown in red (black). The inter-population coupling strengths are equal for all value of values of ΔΩ. Hence, the curve in Fig. 9B3 corresponds to 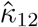 and 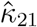. The decoupled solution goes unstable at a torus bifurcation (blue stars) ΔΩ ≈ −0.3 and restabilizes at a second torus bifurcation at ΔΩ ≈ 0.3. The frequency-locked solution branches off the decoupled solution at a pitchfork bifurcation at ΔΩ ≈ −0.15 and ΔΩ ≈ 0.15. As in the *q* = 0.5 case, for the frequency-locked solution, the two populations have non-zero within-population synchrony and intra-/inter-population coupling strengths. However, the within-population synchrony and the intra-population coupling strength are non-longer equal (*ρ*_1_ ≠ *ρ*_2_, 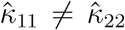). We note that the trivial solution (*ρ_μ_* = 0, 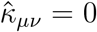), which is stable for all values of ΔΩ, is not shown in Fig. 9B, as it would have obscured the unstable region of the decoupled solution.

Finally, we performed a two-parameter continuation in ΔΩ and *q* (Fig. 9C). The saddle-node curves, which demarcate the region of existence for the coupled solution, are shown in green. The coupled solution exists between the two green curves. The orange curve corresponds to the pitchfork bifurcation, where the frequency-locked solution branches off the decoupled solution. There is a global bifurcation at *q* = 0.2 = 2Δ/λ and *q* = 0.8 = 8Δ/λ, when the decoupled solution ceases to exist and, as such, there is no longer a pitchfork bifurcation. The torus bifurcation curve, where the decoupled solution changes stability, is plotted in blue. In the region between the torus bifurcation curve and the saddle-node bifurcation curve, the two populations have non-trivial dynamics. The inter-population coupling is weak enough that the populations do not entrain. Hence, the mean phases precess at different rates, and as such, the phase difference oscillates in time. As a result, the coupling strengths oscillate as the populations move in- and out-of-phase with each other. Given that the coupling strengths and the synchrony are interdependent, the synchrony variables also oscillate in time.

## 5 Describing the full adaptive network using coupled populations evolving according to mean-field dynamics

In this section, we aim to approximate a network of Kuramoto oscillators (equation (4)) with adaptivity given by symmetric PDDP (equation (3) with *φ* = 0) using two coupled populations evolving according to the mean-field dynamics. As shown in Section 2.2, the full adaptive network with symmetric PDDP is itself a good approximation of the full adaptive network with symmetric STDP. Here, we optimise parameters of the coupled mean-field equations to best approximate the full network.

### 5.1 Optimising the parameters of the two-population mean-field to approximate the full adaptive network

To approximate the full adaptive network, we consider the two-population mean-field model

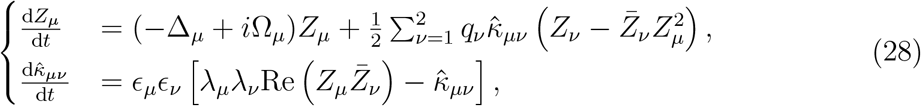

where *μ, ν* ∈ {1, 2} are population indices, *Z_μ_* is the order parameter of population *μ*, and 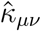 is the average weight from population *ν* to population *μ*. Synthetic data from the full adaptive network is generated by simulating a network of *N* = 100 Kuramoto oscillators (equation (4)) with adaptivity given by symmetric PDDP (equation (3) with *φ* = 0). Further details on the generation of synthetic data, as well as descriptions of the test and training sets can be found in Supplemental Material B.

To describe the full adaptive network using the two-population mean-field approximation (28), we optimise *∈_μ_* and λ*_μ_* with *μ* ∈ {1, 2}, as well as the proportion of oscillators allocated to the first population denoted by *q*_1_. Note that adaptivity parameters within and between populations (λ_1_, λ_2_, *∈*_1_, *∈*_2_) are not the same as adaptivity parameters between oscillators in the full network (λ and *ε*). It is also unclear what is the optimal proportion of oscillators to allocate to each population, we therefore optimise the allocation of oscillators as follows. We sort oscillators by the average value of their mean outgoing coupling (i.e. 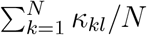), and allocate the first 100 × *q*_1_% to the first population, and the rest to the second population (*q*_2_ = 1 – *q*_1_). This results in populations of size *N*_1_ and *N*_2_, and initial conditions for equation (28) can be obtained from the initial conditions used to simulate the full adaptive network. For each initial condition, the parameters Ω and Δ are obtained as the median frequency and half of the interquartile range of the frequency of oscillators in each population, respectively. The optimisation is performed as a sweep over *q*_1_ = {0.1, 0.2, 0.3, …, 0.9}, and for each value of *q*_1_, 36 local optimisations over the remaining parameters are carried out. Each local optimisation starts from a random set of parameters, and consists in successive optimisations using patternsearch and fminsearch (MatlabR2021a) over all initial conditions in the training set.

The cost function minimised by the optimisation is the average over initial conditions in the training set of

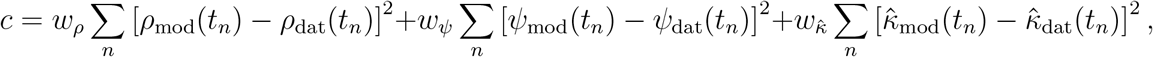

where *t_n_* are all the time points corresponding to the second half of the simulation (to limit the influence of transients), *ρ* is the modulus of the order parameter of the entire system, *ψ* is the (unwrapped) phase of the order parameter of the entire system, 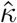 is the mean coupling weight of the entire system, and subscripts “mod” and “dat” refer to equation (28) and synthetic data from the full adaptive network, respectively. The coefficients *ω_ρ_, ω_ψ_*, 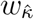 are chosen to ensure that the costs corresponding to *ρ*, *ψ*, and 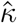 are on a similar scale. For the two-population mean-field approximation, the order parameter of the whole system is obtained as *Z* = *q*_1_*Z*_1_ + *q*_2_*Z*_2_, and similarly the average coupling is obtained as 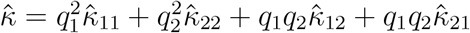.

### 5.2 Performance on training set and test set

The optimised two-population mean-field approximation with the lowest cost was found for *q*_1_ = 0.6 (dark blue in Fig. 10), and its parameters are given in Table C.1 of Supplemental Material C. To test the model performance, we construct three different control models; (i) a constant model defined from initial conditions by *ρ* = *ρ*_0_ and 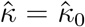, (ii) a two-population mean-field approximation with *q*_1_ = 0.6 as in the fully optimised model, but without adaptivity (*∊*_1_ = *∊*_2_ = 0), and (iii) a two-population mean-field approximation where λ_1_λ_2_ = 25 and *∊*_1_*∊*_2_ = 0.5 are chosen to match *λ* and *ϵ*, respectively. For the third control model, we performed as sweep over *q*_1_ and found the best fit for *q*_1_ = 0.95.

**Figure 10:**
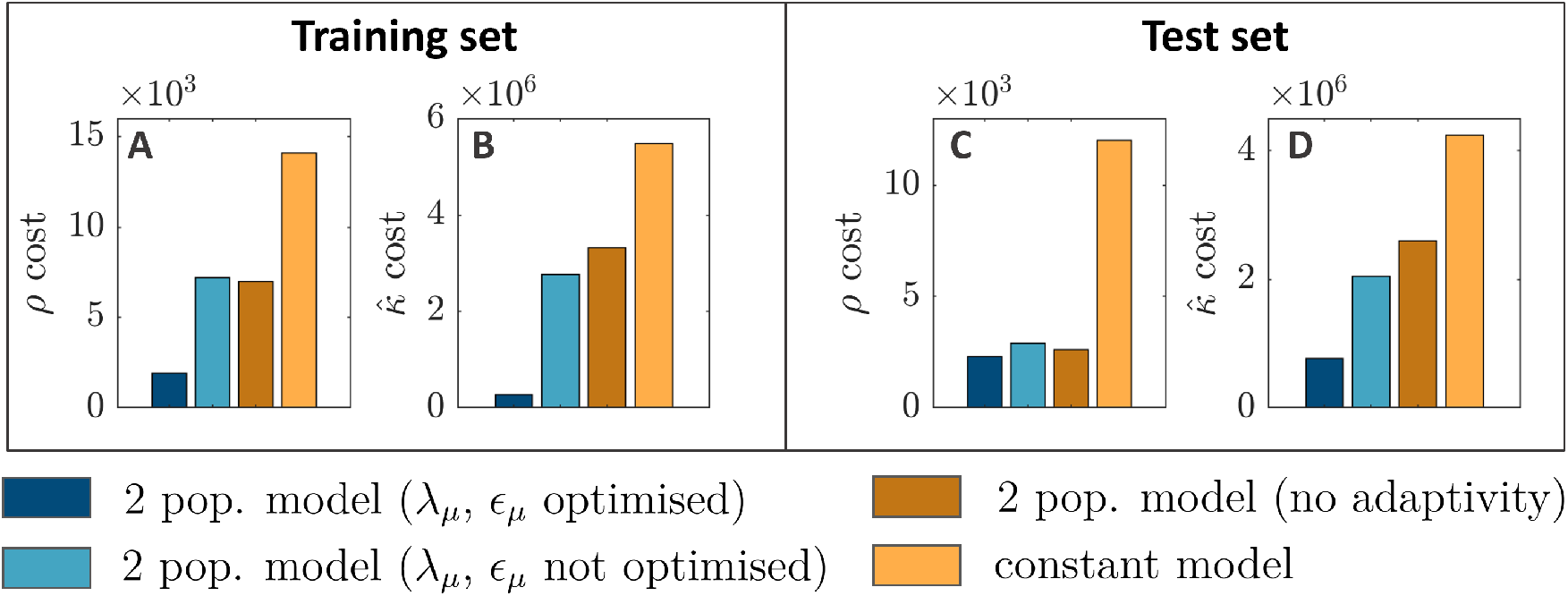
Performance of the two-population mean-field approximation. Each panel compares the sum of squared differences over time between model and synthetic data for *ρ* or 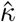, averaged over the training set or test set. The two-population mean-field approximation with optimised plasticity parameters is shown in dark blue, the two-population mean-field approximation with plasticity parameters obtained from the full system is shown in light blue, the two-population model without adaptivity is shown in dark orange, and the constant model is shown in orange. Training and test sets are composed of 13 and 20 trajectories, respectively, with varied initial conditions (more details in Section B in the Appendix).

The fully optimised two-population mean-field approximation with the lowest cost on the training set is better at describing the dynamics of *ρ* and 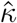 both on the training set and on the test set than the controls shown in Fig. 10. This shows that to best reproduce synthetic data from the full adaptive model, it is necessary to consider both the evolution of the order parameter and of the average weights, and to optimise the within/between population adaptivity parameters. The constant model is shown in light orange, the two-population mean-field approximation with no adaptivity is shown in dark orange, and the two-population mean-field approximation where the plasticity parameters are not optimised is in light blue. In the test set, the greatest improvement of the optimised two-population mean-field approximation over the controls is for 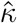 (Fig. 10D). With the exception of the constant model, the controls already approximate *ρ* well in the test set (Fig. 10C). Representative trajectories of the two-population mean-field approximation obtained from initial conditions in the test set approximate the network synchrony, phase, and average coupling of synthetic data from the full adaptive network (see Figure C.10 in the Appendix). The approximation of 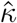 is overall better when *∊_μ_* and λ*_μ_* are optimised. We note that while transients were not included in the cost function used to fit to synthetic data, they are relatively well described by the model when starting from random connectivity with different means in the test set (see Fig C.10).

## 6 Discussion

Using simulations, we showed that PDDP and ebPDDP can provide useful approximations of STDP. In particular, single-harmonic rules can approximate simple forms of symmetric STDP, while multi-harmonic rules are required to accurately approximate causal STDP. In the latter case, the accuracy of the approximation increases with the number of Fourier coefficients before reaching a plateau. The Fourier coefficients can be easily computed using analytical expressions only involving causal STDP parameters (equation (9)). We found ebPDDP to be a slightly better approximation than PDDP both in the case of symmetric STDP and causal STDP. This is expected since contrary to PDDP, ebPDDP restricts synaptic weight updates to spiking events, which is conceptually closer to STDP. One limitation of approximating STDP using plasticity rules based on phase difference is that the evolution of phase differences is required to be slower than the phase dynamics (Lücken et al., 2016). Under this assumption, the intricate evolution of coupling weights can be well approximated by PDDP and ebPDDP (see e.g Fig. C.1 and Fig. C.5 in Supplemental Material C). For best results, the STDP function should also decay to zero faster than the network mean frequency for both positive and negative spike-timing differences (in the case of causal STDP to avoid both components in equation (6) strongly overlapping).

We derived exact expressions for the evolution of the average coupling weight in networks of phase oscillators with PDDP and ebPDDP. These expressions make no assumption about the underlying network, and are compatible with any plasticity rule based on phase difference as long as it can be expended as a Fourier series of the phase difference. As a proof of principle, we focussed on mean-field approximations based on the average coupling evolution for two-cluster states in a population of adaptive Kuramoto oscillators, and performed a bifurcation analysis to highlight the different possible behaviours. Despite its simplicity, the Kuramoto model is a prototypical model to study synchronisation in neuroscience, with diverse applications including whole-brain modelling, epilepsy, and Parkinson’s disease (Cumin & Unsworth,2007; Breakspear, Heitmann, & Daffertshofer, 2010; Cabral et al., 2014; Schmidt, LaFleur, de Reus, van den Berg, & van den Heuvel, 2015; Ponce-Alvarez et al., 2015; Finger et al., 2016; Asllani, Expert, & Carletti, 2018; Weerasinghe et al., 2019; Bick et al., 2020; Weerasinghe et al., 2021; Duchet, Sermon, Weerasinghe, Denison, & Bogacz, 2022).

Our framework could easily be adapted to consider more biologically realistic oscillator models, such as the *θ*-neuron model or the formally equivalent quadratic integrate-and-fire (QIF) neuron model. Like the Kuramoto model, the *θ*-neuron model is amenable to the Ott-Antonsen ansatz, and as such, is amenable to exact mean-field description (Luke, Barreto, & So, 2013) (for the QIF model use the equivalent Lorentzian ansatz (Montbrió,Pazó, & Roxin, 2015)). The model analysis would be identical, but given the explicit phase dependency in the coupling, we can expect a richer set of dynamics for a network of *θ*-neurons than observed here for Kuramoto oscillators. We could also include explicit synaptic variables, as in (Byrne, O’Dea, Forrester, Ross, & Coombes, 2020), to more accurately model synaptic processing. More generally, even detailed neuron models, such as the Hodgkin-Huxley model, can be approximated by networks of phase oscillators through phase reduction (Brown, Moehlis, & Holmes, 2004; Pietras & Daffertshofer, 2019). Although the Ott-Antonsen ansatz may not be appropriate for such networks, the validity of using PDDP in place of STDP rules holds and alternative reduction techniques could be used to construct a mean-field approximation. Previous studies of adaptive oscillators (Berner et al., 2019), and our full network simulations, point to the existence of three-, four- and five-cluster states. Although computationally expensive and, perhaps, numerically challenging, both the bifurcation analysis and model fitting could be extended to consider more than two clusters.

Combining theoretical insight and data-driven inference, we fitted a two-cluster meanfield approximation to a full adaptive network of Kuramoto oscillators in order to obtain a low-dimensional representation of the full system. While the two-cluster mean-field model can approximate the full adaptive network on the test set, the accuracy of the approximation could be improved in several ways. First, a richer training set could be used. Second, the mean-field description of the order parameter is not exact, and a moment or cumulant approach may capture more of the full system dynamics (Tyulkina, Goldobin, Klimenko, & Pikovsky, 2018). Third, considering more than two populations may provide a more accurate approximation. Fourth, rather than allocating oscillators to clusters by thresholding their mean outgoing coupling, a finer partition could be learnt (Snyder, Zlotnik, & Lokhov, 2020). Nevertheless, data-driven inference of low-dimensional representations of phase-oscillator networks (Thiem, Kooshkbaghi, Bertalan, Laing, & Kevrekidis, 2020; Snyder et al., 2020; Fialkowski et al., 2022) is a promising approach to approximate the behavior of networks with STDP. Once clusters have been identified, a very recent mean-field technique based on the collective coordinate method can be used (Fialkowski et al., 2022).

More general types of STDP call for extensions of our framework. First, one would naturally expect that the strength of plastic connections are bounded. We included soft bounds in our investigation of symmetric STDP through a dampening term (see equation (3)), which is conserved in the average coupling (equation (13)), and is therefore present in the mean-field analyses carried out in Section 4 and 5. While neither soft nor hard bounds can easily be included in the average coupling derivation for causal STDP, we show that short transients of causal STDP with hard bounds are well captured by causal PPDP/ebPPD without bounds (see Fig. C.6 and Fig. C.7 in Supplemental Material C). Second, we primarily focused on symmetric STDP rules, that is, interchanging the order of spikes does will have little to no effect on the change of connection strength. For such adaptation, the network naturally forms clusters of strongly connected units (cf. Fig. 3). By contrast, the causality of traditional asymmetric STDP rules will be reflected in the network structure as shown in Fig. 5 and highlighted very recently in (Thiele, Berner,Tass, Schöll, & Yanchuk, 2023). To analyze the mean-field dynamics of such networks, a natural approach would be to computationally identify emerging feed-forward structures in such networks instead of looking for clusters. In this case, finding a corresponding lowdimensional description as in Section 4 is more challenging. Third, we consider STDP adaptation rules that depend on pairs of oscillator states. More elaborate, spike-based rules such as triplet interactions have recently attracted attention (Pfister & Gerstner, 2006; Montangie, Miehl, & Gjorgjieva, 2020). It would be interesting to have such adaptation reflected in the STDP model as these could be interpreted as “higher-order” network effects (cf. (Bick, Gross, Harrington, & Schaub, 2021)).

As a step towards low-dimensional description of adaptive networks with STDP, our framework has implications for the study of long-term neural processes. In particular, the effects of clinically available DBS quickly disappear when stimulation is turned off, thus stimulation needs to be provided continuously. To spare physiological activity as much as possible, it would therefore be highly desirable to design stimuli aimed at eliciting long-lasting effects. Continuous stimulation can also lead to habituation, where stimulation benefits diminish considerably over the years in some patients with essential tremor (Fasano & Helmich, 2019). Optimising brain stimulation to have long-lasting effects so far relied on computational studies e.g. (Tass & Majtanik, 2006; Popovych & Tass, 2012; Ebert, Hauptmann, & Tass, 2014; Popovych et al., 2015; Manos, Zeitler, & Tass, 2018), or on analytical insights under the simplifying assumption that spiking is only triggered by stimulation pulses (Kromer & Tass, 2020). Other analytical approaches based on mean-field models are focused on short-term changes due to stimulation and do not consider plastic changes (Duchet et al., 2020; Weerasinghe et al., 2019, 2021). Our framework offers an alternative, where exact evolution equations for the average coupling within and between neural populations could inform the development of new therapies aimed at maximising the long-term effects of brain stimulation.

## Acknowledgements

We are grateful to Rafal Bogacz and Alessandro Torcini for helpful discussions. The authors would like to acknowledge the use of the University of Oxford Advanced Research Computing (ARC) facility in carrying out this work http://dx.doi.org/10.5281/zenodo.22558

## Data availability statement

No data were collected as part of this work.

## Supplemental Material

## A Simulation methods used to compare STDP to PDDP

Using simulations, we compare networks of Kuramoto oscillators with adaptivity given by symmetric or causal STDP, to the same networks with adaptivity given by the corresponding single- or multi-harmonic PDDP and ebPDDP rules.

### A.1 Symmetric learning rules

We simulate networks of *N* = 60 Kuramoto oscillators evolving according to equation (4), where the natural frequencies *ω_k_* are sampled from a normal distribution of mean Ω = 10*π* (5Hz) unless otherwise stated and standard deviation Δ. A normal distribution was chosen over a Lorentzian distribution to avoid extreme natural frequencies which lead to increased variability in simulation repeats. Initial conditions are also sampled from normal distributions. Phases *θ_k_*(*t* = 0s) are sampled from 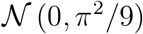, and couplings *κ_kl_*(*t* = 0s) from 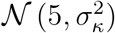. The evolution of the coupling weights *κ_kl_* is simulated according to equation (2) (symmetric STDP) together with the weight decay 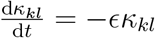, equation (3) with *φ* = 0 (symmetric PDDP), or equation (5) (symmetric ebPDDP). For ebPDDP, we use 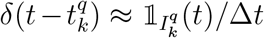 where 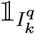 is the indicator function of 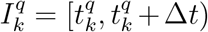 and Δ*t* is the simulation time step. The network is simulated for 150s, using the Euler method with Δ*t* = 1ms unless otherwise stated. We use *ϵ* = 0.5, and the values of other parameters are detailed in Fig. 3 and Fig. C.1.

### A.2 Causal learning rules

Error metrics (described below) are computed between multi-harmonic PDDP or ebPDDP and causal STDP for selected regions of parameter space. To limit computational cost, we constrain causal STDP parameters based on data and biological motivations. We use *τ_+_* = 16.8ms and *τ*___ − = 33.7ms, which were obtained by fitting equation (1) to experimental data (data published in (Bi & Poo, 1998), and fit in (Bi & Poo, 2001)). Several experimental studies have reported the LTD time constant *τ*___ to be larger than the LTP time constant *τ*_+_, e.g. in the rat hippocampus (Bi & Poo, 1998), somatosensory cortex (Feldman, 2000), and visual cortex (Froemke & Dan, 2002). We take *A*_+_ = 0.2 and *A*_/*A*_+_ = *β*. Since the time window for LTD is twice as long as the time window for LTP, we choose *β* = 0.5 to study a balanced situation where LTP dominates for shorter spike-timing differences and LTD dominates for longer spike-timing differences, and *β* = 1 to study a situation where LTD always dominates.

We simulate networks of Kuramoto oscillators as for symmetric STDP (see previous section) with the following differences. Initial couplings *κ_kl_*(*t* = 0s) are sampled from 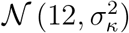. The evolution of the coupling weights *κ_kl_* is simulated according to equation (1) (causal STDP), equation (6) (causal PDDP), or equation (7) (causal ebPDDP). For PDDP and ebPDDP, the Fourier expansion of *F* is truncated after *N_f_* coefficients. No weight decay term is included.

The comparison of multi-harmonic PDDP or ebPDDP to causal STDP relies on four error metrics. In the following error metric definitions, we use the subscript or superscript X to refer to quantities corresponding to multi-harmonic PDDP or ebPDDP. To compare the time evolution of network synchrony we define

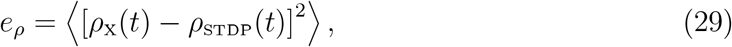

where 〈.〉 denotes time averaging over the duration of the simulation [0*, *t**_max_], and *ρ*(*t*) = |*Z*(*t*) | is the network synchrony. To compare the time evolution of the network average coupling we compute

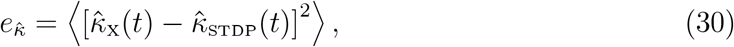

where the average coupling is given by equation (10). The time evolution of the weight distribution is compared using

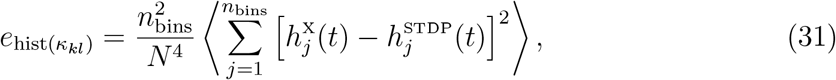

where *h_j_* (*t*) is the count of couplings weights falling into the *j*^th^ bin in the histogram of coupling weights at time *t* (bin boundaries are taken identical across rules and time for a given set of parameters). The fourth error metric 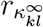 is the Pearson’s correlation coefficient between the STDP coupling matrix and the PDDP/ebPDDP coupling matrix at the last stimulation time point. The scaling factors in equation (31) and the choice of a correlation measure for the fourth metric ensure that the corresponding metrics are independent of the size of the network, and of the number of bins. For each set of parameters, the four metrics are averaged across five repeats. For a given repeat and a given set of parameters, the same random samples are used as initial conditions to compare the network evolution between plasticity rules.

## B

### Methodological details pertaining to the optimisation of the two-population mean-field approximation

Synthetic data is generated by stimulating *N* = 100 Kuramoto oscillators evolving according to equation (4), with adaptivity given by symmetric PDDP (equation (3) with *φ* = 0, λ = 25, and *ϵ* = 0.5). Oscillator natural frequencies *ω_k_* are sampled from a Lorentzian distribution of center Ω = 10*π* (5Hz) and width Δ = 0.6*π*. Initial phases are sampled from a Von Mises distribution of standard deviation *π*/4. Initial couplings *κ_kl_*(*t* = 0s) are sampled from 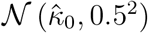. The network is simulated for 20s, using the Euler method with Δ*t* = 0.1ms.

We create a training set and a test set based on different initial conditions and in particular various 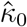. The training set corresponds to 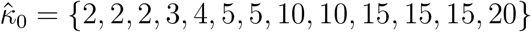, and the test set to 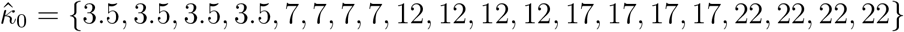. For each trajectory in the training and test sets, natural frequencies, initial phases, and initial couplings are sampled from their respective distributions. Repeated 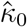 values therefore correspond to different systems with different initial synchrony.

For optimisation speed and accuracy the two-population mean-field approximation is simulated using the variable order solver ode113 in Matlab (variable-step, variable-order Adams-Bashforth-Moulton solver of orders 1 to 13).

## C Supplementary figures and tables

**Table C.1:** Best parameters of two-population mean-field approximation. Parameters correspond to equation (28). Full network parameters used to generate synthetic data for the optimisation are given in Section B.

| Parameter | $\epsilon_1$ | $\epsilon_2$ | $\lambda_1$ | $\lambda_2$ |
| --- | --- | --- | --- | --- |
| Value | 0.0263 | 3.0262 | 5.2985 | 5.4759 |

---

**Figure C.1:**
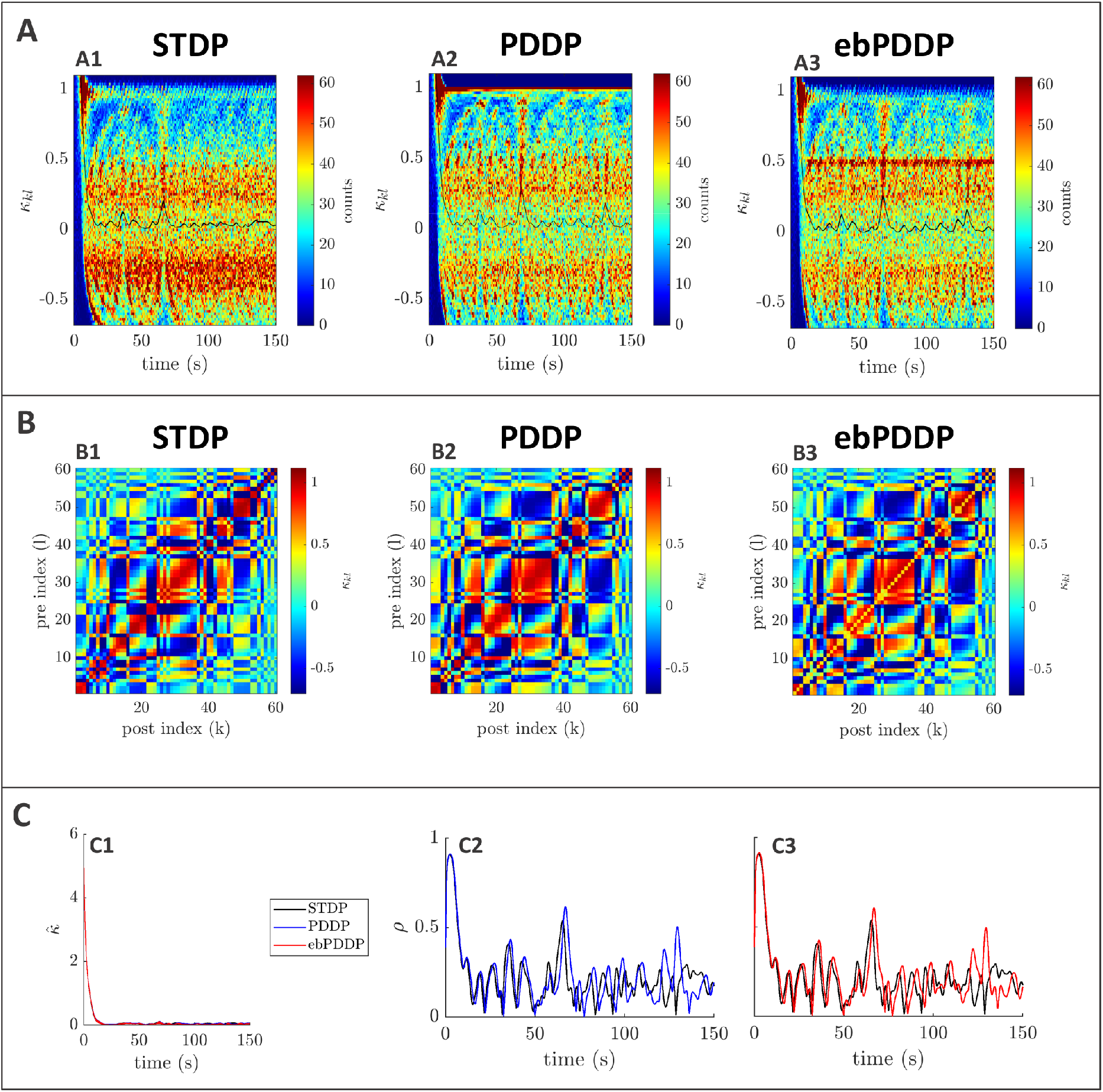
Comparison between symmetric STDP and PDDP in a Kuramoto network (desynchronised state). **A:** Evolution of the distribution of coupling weights with time (100 bins at each time point). The average weight is represented by a thin black line. STDP is shown in A1, PDDP in A2, and ebPDDP in A3. **B:** Coupling matrix at *t* = 150s, with oscillators sorted by natural frequency. STDP is shown in B1, PDDP in B2, and ebPDDP in B3. **C:** Time evolution of average coupling (C1, error bars represent the standard error of the mean over 5 repeats, too small to see) and network synchrony (C2-C3). STDP is shown in black, PDDP in blue, and ebPDDP in red. [*a* = 0.025822, *b* = 0.049415, *σ_κ_* = 3, Δ = 0.2*π*, Ω = 10*π* (5Hz)]

**Figure C.2:**
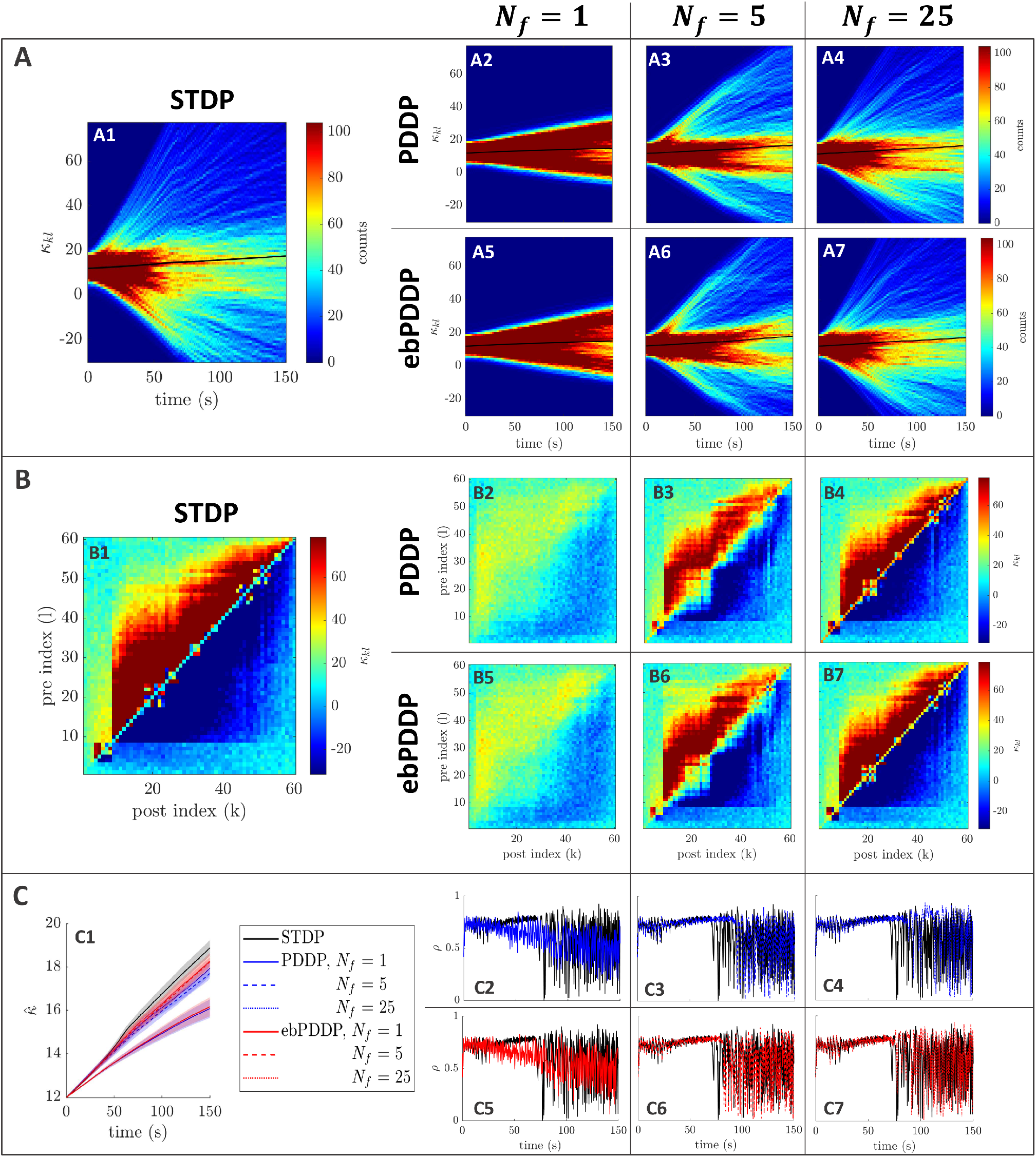
Comparison between STDP and PDDP in a Kuramoto network, [*β* = 0.5, *σ_κ_* = 3, Δ = 1.8*π*, Ω = 10*π* (5Hz)]. Results for PDDP and ebPDDP are shown for 1, 5, and 25 Fourier components *N_f_* (first, second, and third column on the right hand side of the figure, respectively). **A:** Evolution of the distribution of coupling weights with time (100 bins at each time point). The average weight is represented by a thin black line. STDP is shown in A1, PDDP in A2-A4, and ebPDDP in A5-A7. **B:** Coupling matrix at *t* = 150s, with oscillators sorted by natural frequency. STDP is shown in B1, PDDP in B2-B4, and ebPDDP in B5-B7. **C:** Time evolution of average coupling (C1, error bars represent the standard error of the mean over 5 repeats) and network synchrony (C2-C7). STDP is shown in black, PDDP in blue, and ebPDDP in red.

**Figure C.3:**
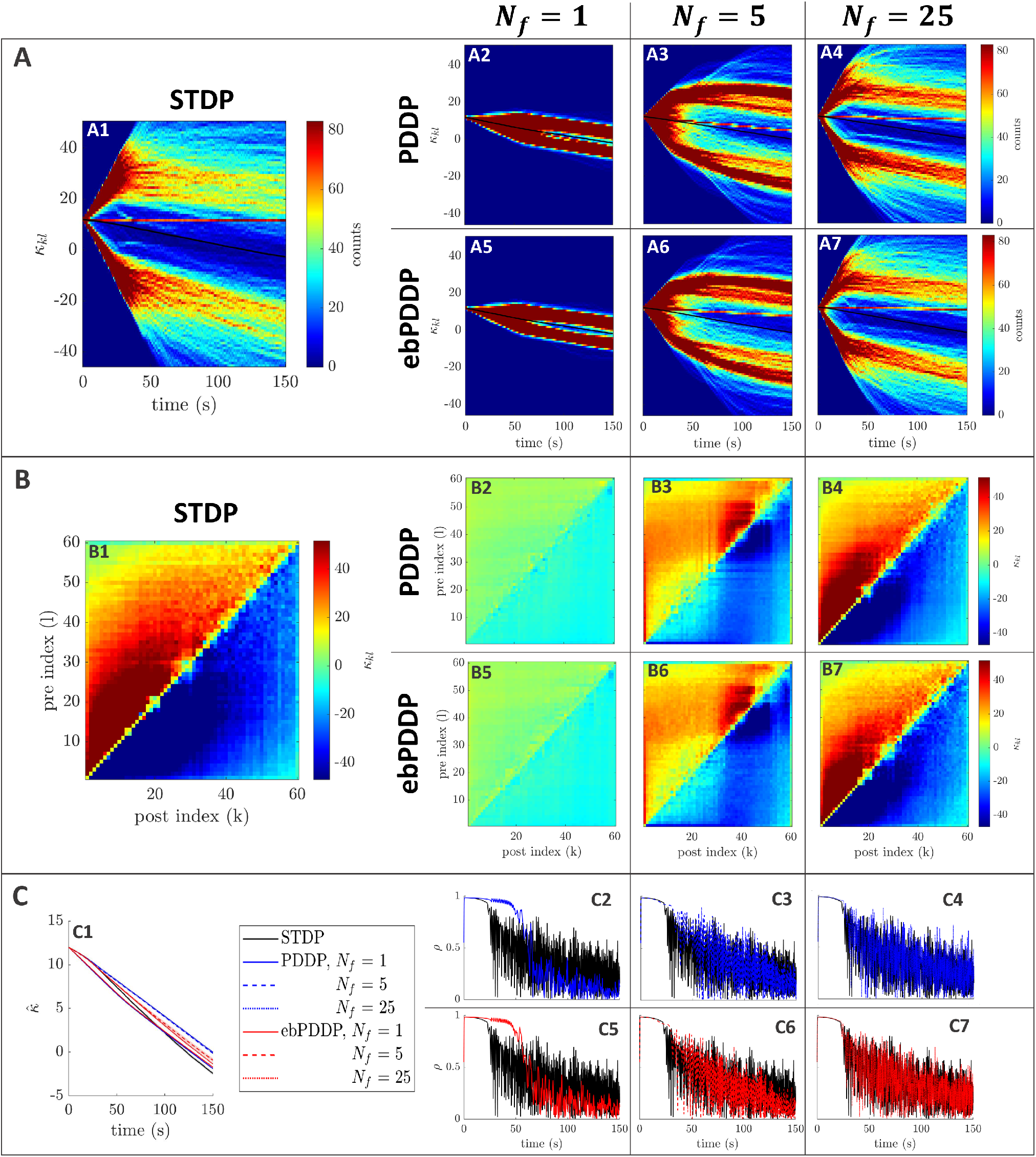
Comparison between STDP and PDDP in a Kuramoto network, [*β* = 1, *σ_κ_* = 0.2, Δ = 0.6*π*, Ω = 10*π* (5Hz)]. Results for PDDP and ebPDDP are shown for 1, 5, and 25 Fourier components *N_f_* (first, second, and third column on the right hand side of the figure, respectively). **A:** Evolution of the distribution of coupling weights with time (100 bins at each time point). The average weight is represented by a thin black line. STDP is shown in A1, PDDP in A2-A4, and ebPDDP in A5-A7. **B:** Coupling matrix at *t* = 150s, with oscillators sorted by natural frequency. STDP is shown in B1, PDDP in B2-B4, and ebPDDP in B5-B7. **C:** Time evolution of average coupling (C1, error bars represent the standard error of the mean over 5 repeats) and network synchrony (C2-C7). STDP is shown in black, PDDP in blue, and ebPDDP in red.

**Figure C.4:**
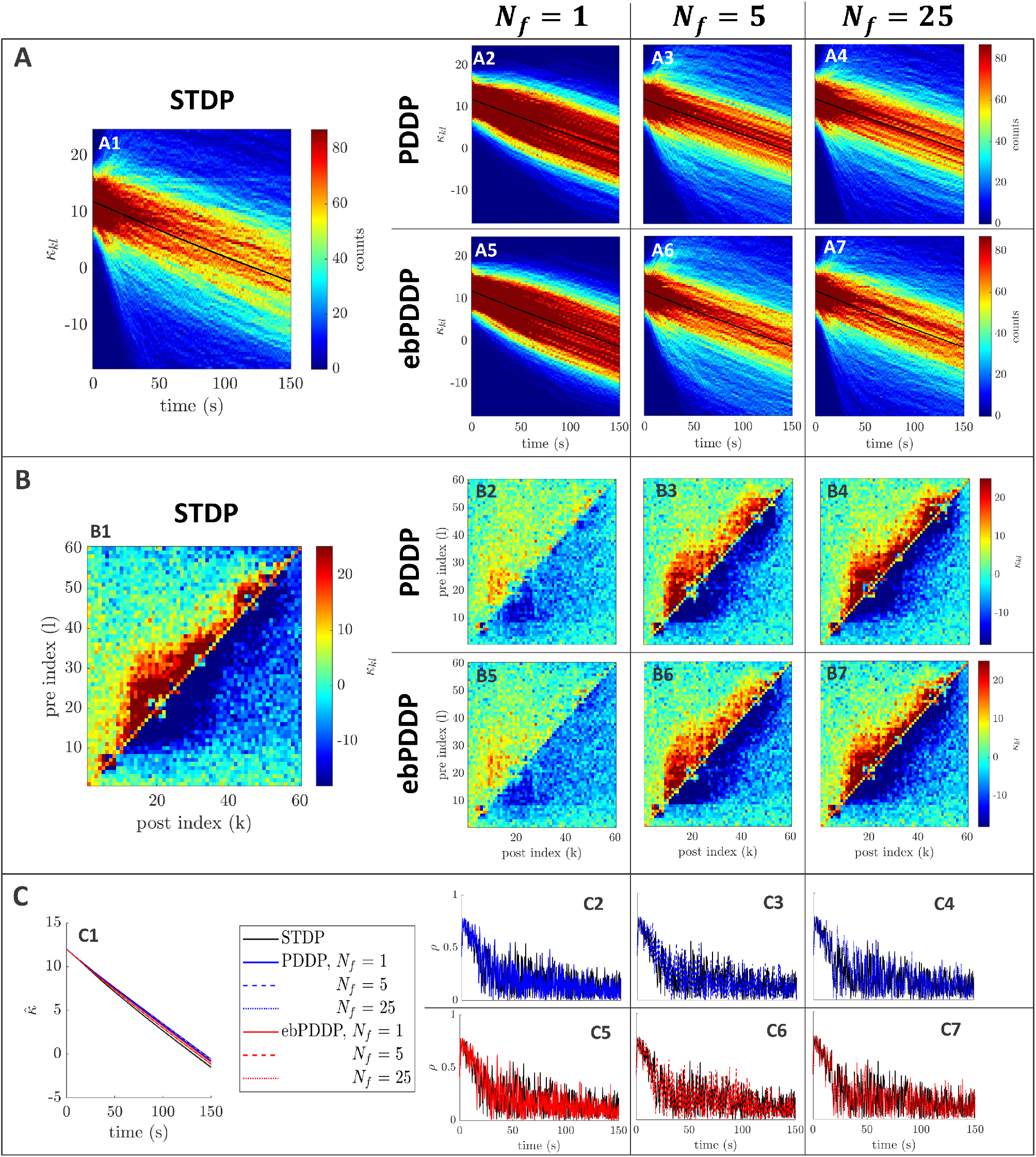
Comparison between STDP and PDDP in a Kuramoto network, [*β* = 1, *σ_κ_* = 3, Δ = 1.8*π*, Ω = 10*π* (5Hz)]. Results for PDDP and ebPDDP are shown for 1, 5, and 25 Fourier components *N_f_* (first, second, and third column on the right hand side of the figure, respectively). **A:** Evolution of the distribution of coupling weights with time (100 bins at each time point). The average weight is represented by a thin black line. STDP is shown in A1, PDDP in A2-A4, and ebPDDP in A5-A7. **B:** Coupling matrix at *t* = 150s, with oscillators sorted by natural frequency. STDP is shown in B1, PDDP in B2-B4, and ebPDDP in B5-B7. **C:** Time evolution of average coupling (C1, error bars represent the standard error of the mean over 5 repeats) and network synchrony (C2-C7). STDP is shown in black, PDDP in blue, and ebPDDP in red.

**Figure C.5:**
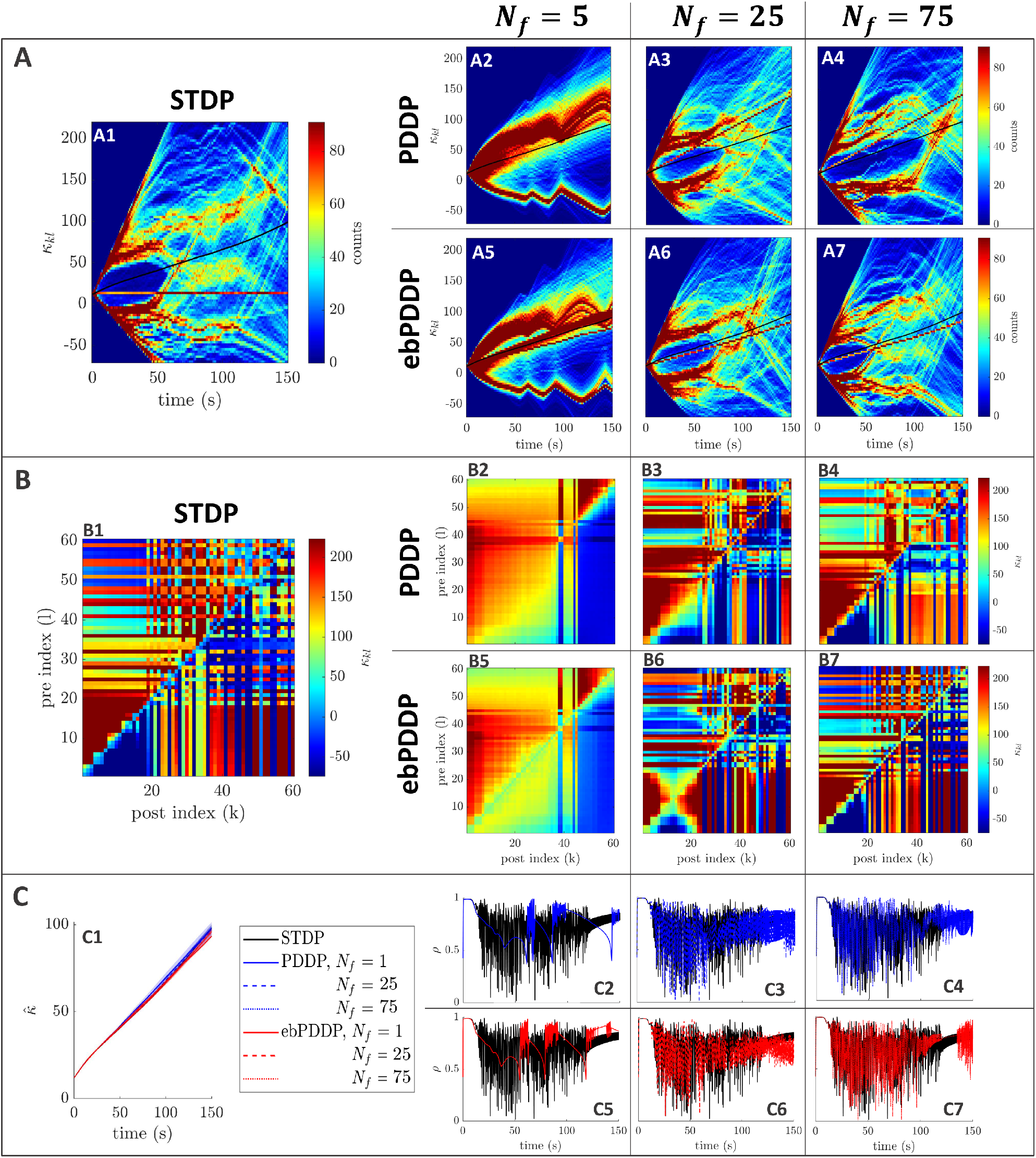
Comparison between STDP and PDDP in a Kuramoto network, [*β* = 0.5, *σ_κ_* = 0.2, Δ = 0.6*π*, Ω = 40*π* (20Hz), Δ*t* = 0.1ms]. Results for PDDP and ebPDDP are shown for 5, 25, and 75 Fourier components *N_f_* (first, second, and third column on the right hand side of the figure, respectively). **A:** Evolution of the distribution of coupling weights with time (100 bins at each time point). The average weight is represented by a thin black line. STDP is shown in A1, PDDP in A2-A4, and ebPDDP in A5-A7. **B:** Coupling matrix at *t* = 150s, with oscillators sorted by natural frequency. STDP is shown in B1, PDDP in B2-B4, and ebPDDP in B5-B7. **C:** Time evolution of average coupling (C1, error bars represent the standard error of the mean over 5 repeats) and network synchrony (C2-C7). STDP is shown in black, PDDP in blue, and ebPDDP in red.

**Figure C.6:**
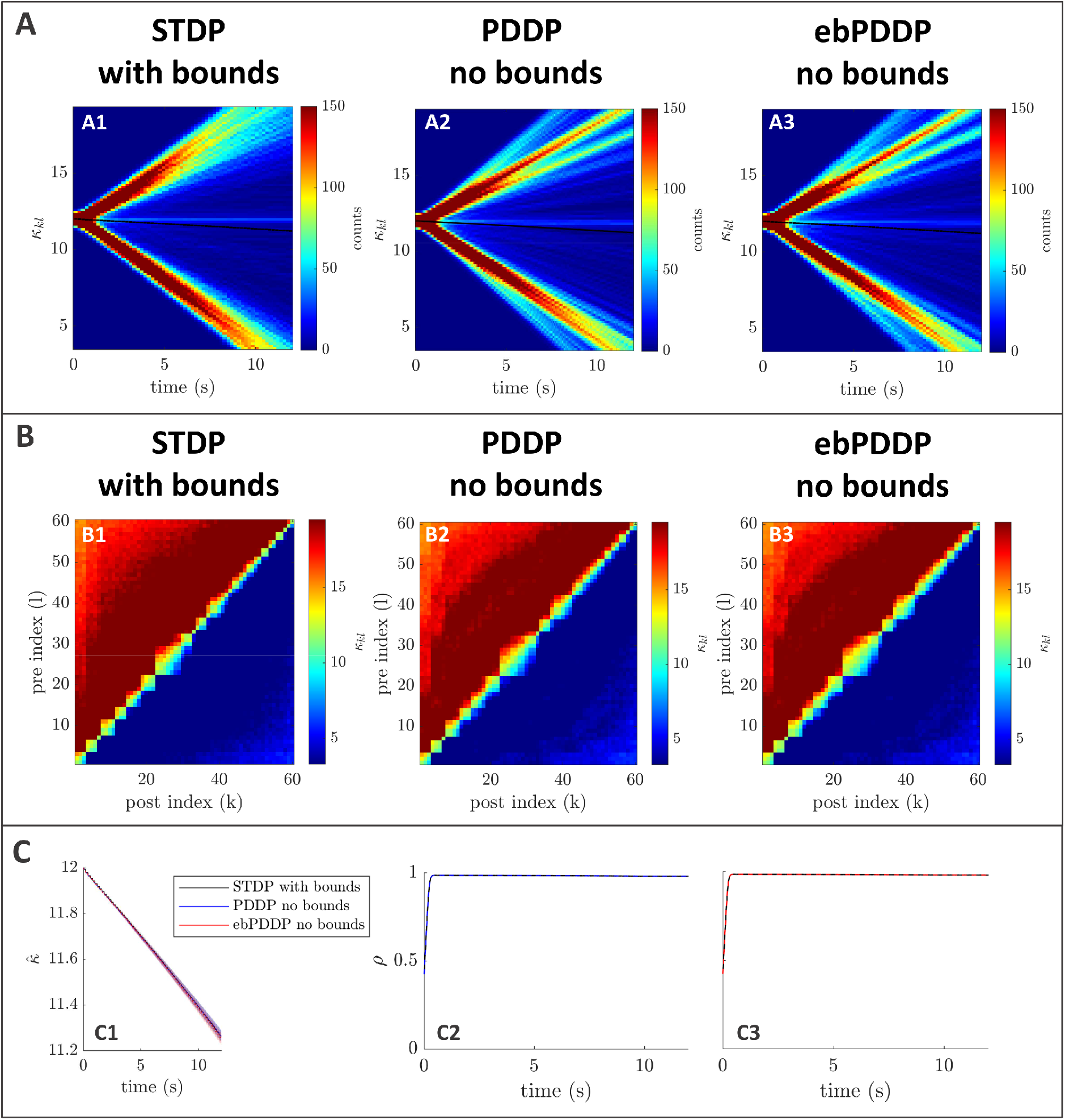
Comparison between STDP with hard bounds and PDDP without bounds in a Kuramoto network, [*β* = 1, *σ_κ_* = 0.2, Δ = 0.6*π*, Ω = 10*π* (5Hz)]. Results for PDDP and ebPDDP are shown for *N_f_* = 40 Fourier components. For STDP, hard bounds are enforced such that no individual weight can go below 0 or above 30. **A:** Evolution of the distribution of coupling weights with time (100 bins at each time point). The average weight is represented by a thin black line. STDP with hard bounds is shown in A1, PDDP without bounds in A2, and ebPDDP without bounds in A3. **B:** Coupling matrix at *t* = 12s, with oscillators sorted by natural frequency. STDP with hard bounds is shown in B1, PDDP without bounds in B2, and ebPDDP without bounds in B3. **C:** Time evolution of average coupling (C1, error bars represent the standard error of the mean over 5 repeats) and network synchrony (C2-C3). STDP with hard bounds is shown in black, PDDP without bounds in blue, and ebPDDP without bounds in red.

**Figure C.7:**
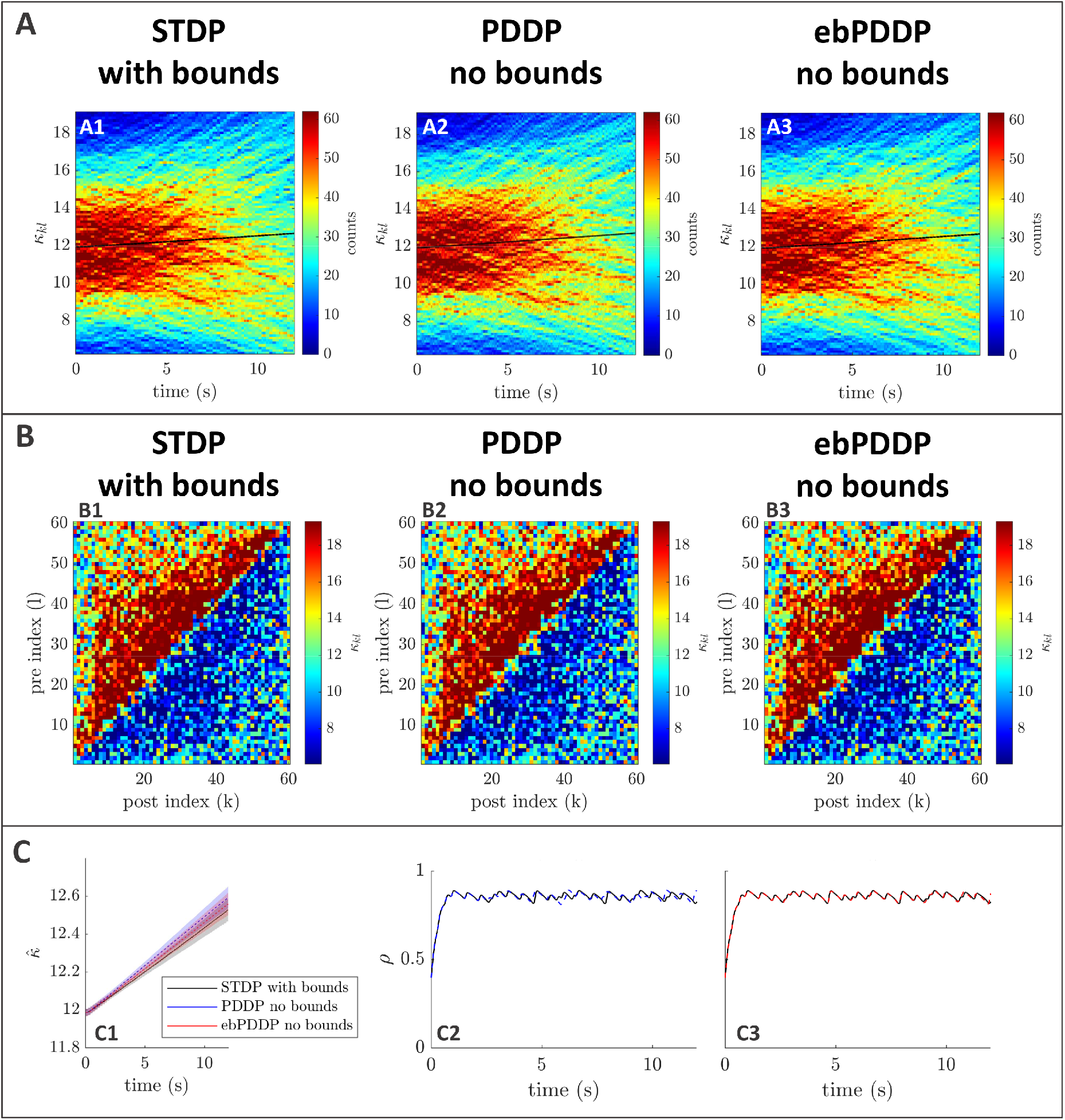
Comparison between STDP with hard bounds and PDDP without bounds in a Kuramoto network, [*β* = 0.5, *σ_κ_* = 3, Δ = 1.8*π*, Ω = 10*π* (5Hz)]. Results for PDDP and ebPDDP are shown for *N_f_* = 40 Fourier components. For STDP, hard bounds are enforced such that no individual weight can go below 0 or above 30. **A:** Evolution of the distribution of coupling weights with time (100 bins at each time point). The average weight is represented by a thin black line. STDP with hard bounds is shown in A1, PDDP without bounds in A2, and ebPDDP without bounds in A3. **B:** Coupling matrix at *t* = 12s, with oscillators sorted by natural frequency. STDP with hard bounds is shown in B1, PDDP without bounds in B2, and ebPDDP without bounds in B3. **C:** Time evolution of average coupling (C1, error bars represent the standard error of the mean over 5 repeats) and network synchrony (C2-C3). STDP with hard bounds is shown in black, PDDP without bounds in blue, and ebPDDP without bounds in red.

**Figure C.8:**
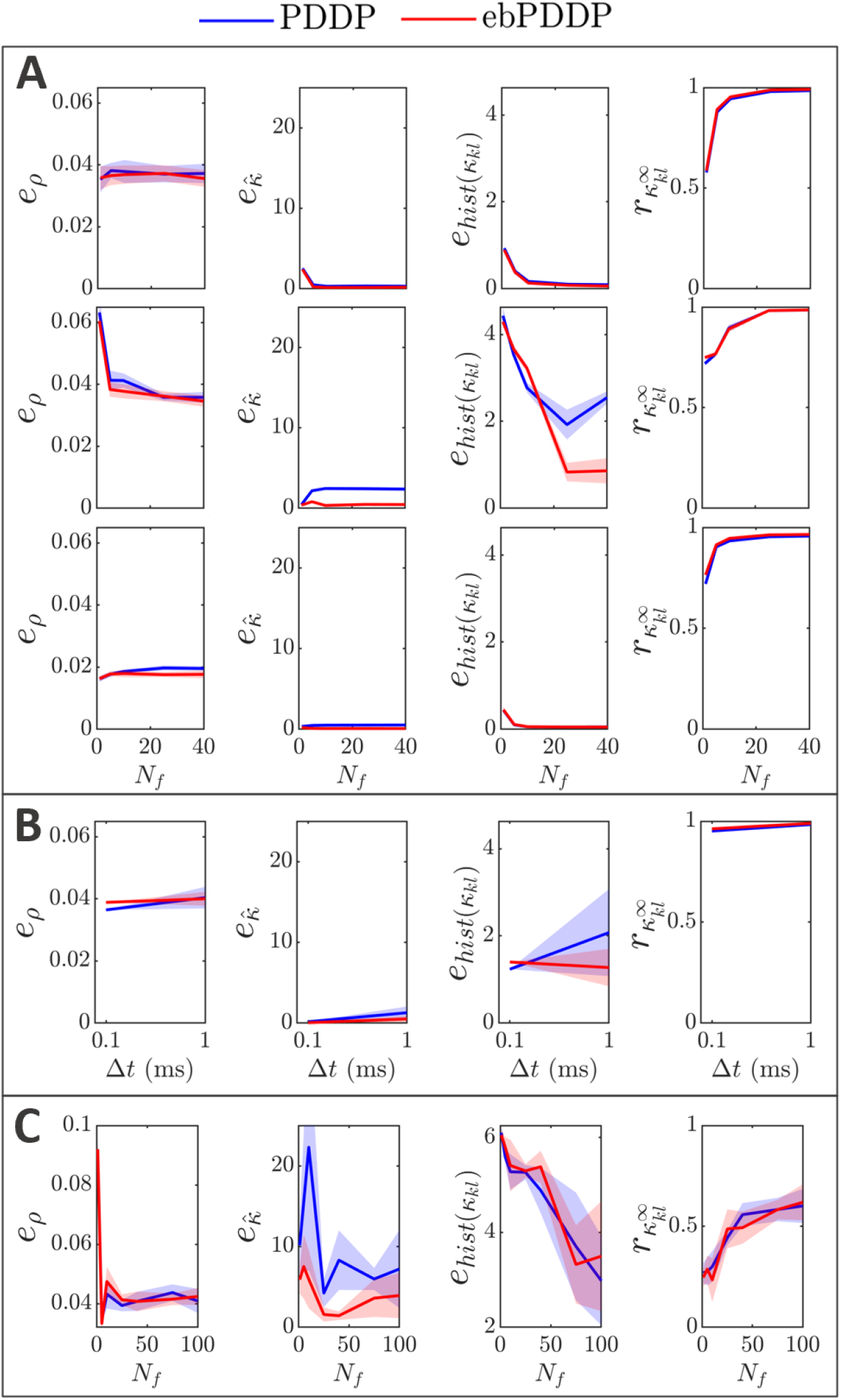
Influence of *N_f_*, Δ*t*, and network frequency on error metrics. Error metrics for PDDP compared to STDP for the time evolution of network synchrony *e_ρ_*, average coupling 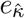 and distribution of weights *e*_hist(*κl*_), and for the coupling matrix at the last stimulation point 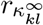 are shown in the first, second, third, and fourth columns, respectively. Results for PDDP are in blue, and for ebPDDP in red. Showing sem error bars over 5 repeats. **A:** Influence of the number of Fourier coefficients *N_f_* on the error metrics for the parameters used in Fig. C.2 (*β* = 0.5, *σ_κ_* = 3, Δ = 1.8*π*, Ω = 10*π*) in the first row, in Fig. C.3 (*β* = 1, *σ_κ_* = 0.2, Δ = 0.6*π*, Ω = 10*π*) in the second row, in Fig. C.4 (*β* = 1, *σ_κ_* = 3, Δ = 1.8*π*, Ω = 10*π*) in the third row. **B:** Influence of the time step Δ*t* for the parameters used in Fig. 5 (*β* = 0.5, *σ_κ_* = 0.2, Δ = 0.6*π*, Ω = 10*π*) and *N_f_* = 40. **C:** Influence of the number of Fourier coefficients *N_f_* on error metrics for the parameters used in the 20Hz example, see Fig. C.5 (*β* = 0.5, *σ_κ_* = 0.2, Δ = 0.6*π*, Ω = 40*π*, Δ*t* = 0.1ms).

**Figure C.9:**
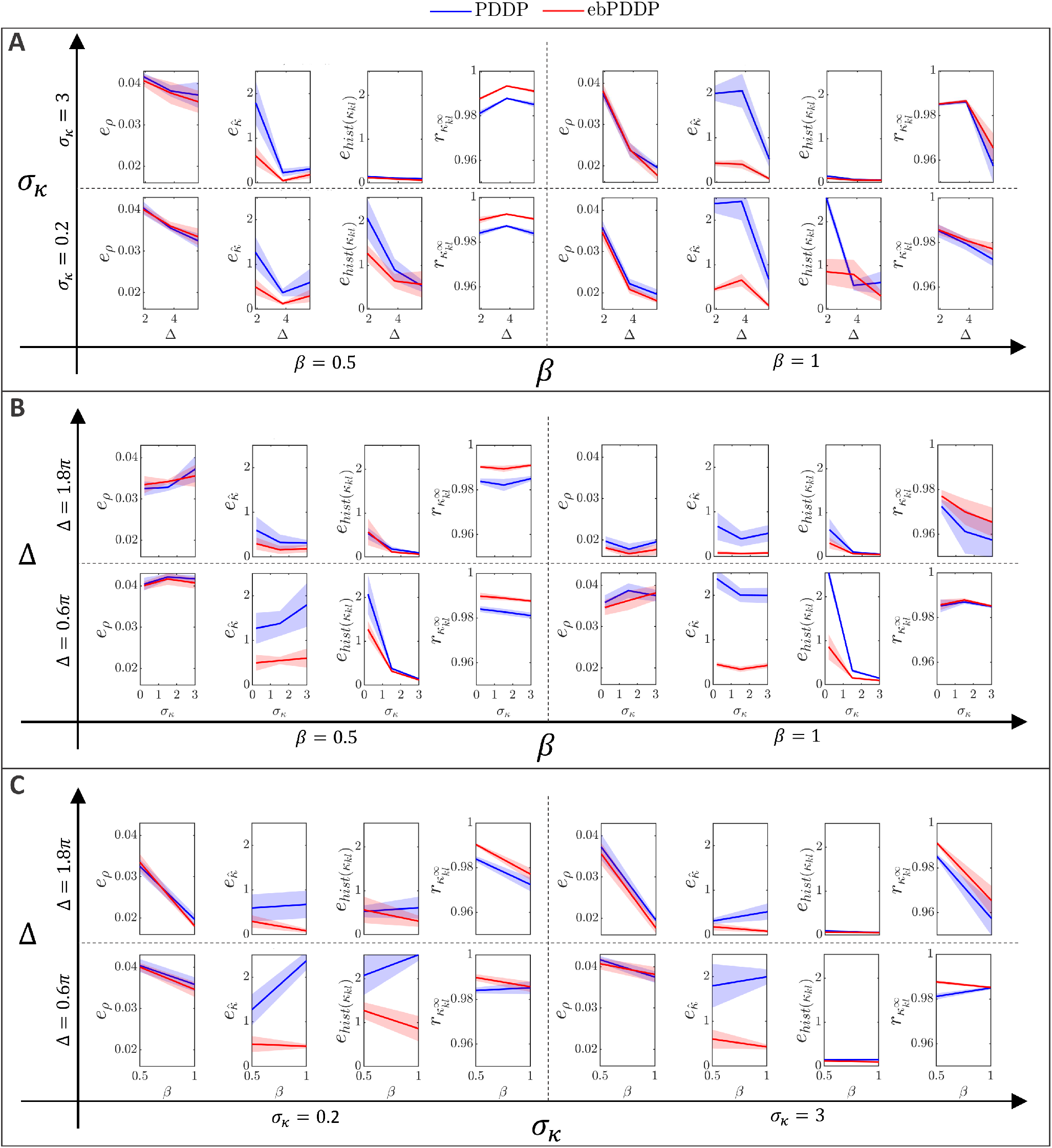
Influence of *σ_κ_*, Δ, and *β* on error metrics (slices through parameter space). Error metrics for PDDP compared to STDP for the time evolution of network synchrony *e_ρ_* (first and fifth columns), average coupling 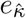 (second and sixth columns) and distribution of weights *e*_*hist*(*κ_kl_*_) (third and seventh columns), and for the coupling matrix at the last stimulation point 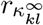 (fourth and eighth columns). Results for PDDP are in blue, and for ebPDDP in red. Each of the panels represents a different slice through parameter space as indicated by the axes. The error metrics are shown for *N_f_* = 40 Fourier components, with sem error bars over 5 repeats.

**Figure C.10:**
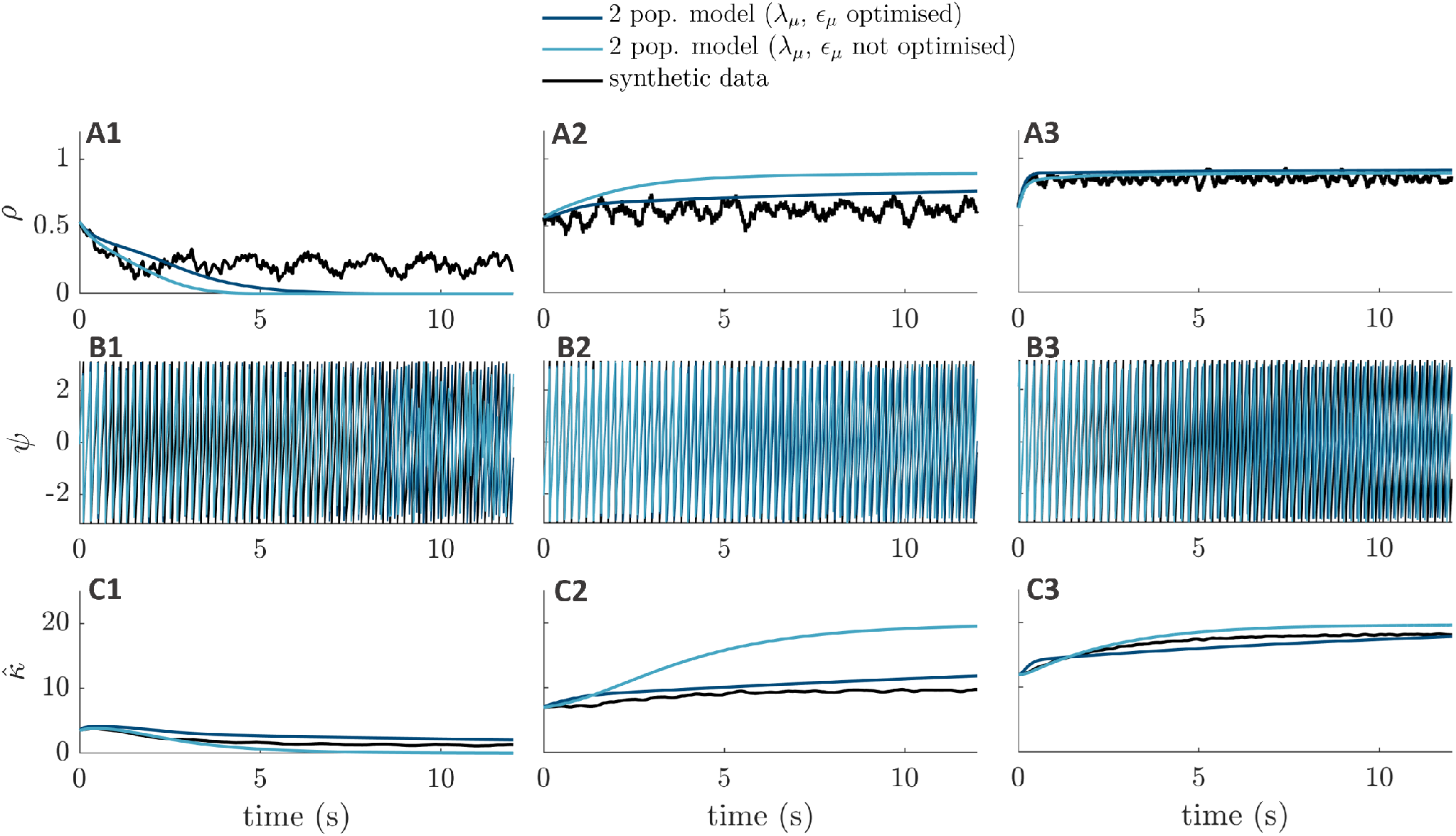
Representative trajectories from initial conditions in the test set. The two-population mean-field approximation with optimised plasticity parameters is shown in dark blue, the two-population mean-field approximation with plasticity parameters obtained from the full system is shown in light blue, and synthetic data from the full adaptive Kuramoto network is shown in black. The first row shows the network synchrony, the second row the network phase, and the third row the network average coupling.

## Footnotes

1 In this work, we call plasticity rules *causal* when temporal precedence is enforced (as in ‘classical’ STDP rules (Bi & Poo, 1998)), and *symmetric* when inverting spike timings/phase differences leads to the same type of adaptation. We stay clear of the ambiguous term *Hebbian* STDP, which is also sometimes used to refer to causal STDP in the experimental literature (Cassenaer & Laurent, 2007; Sgritta et al., 2017.

2 The division by 2 ensures that this rule scales similarly to the STDP rule and the PDDP rule with continuous updates.

3 Note that equilibrium points of the effective dynamics correspond to periodic solutions of the full system.

